# An integrated single-cell reference atlas of the human endometrium

**DOI:** 10.1101/2023.11.03.564728

**Authors:** Magda Marečková, Luz Garcia-Alonso, Marie Moullet, Valentina Lorenzi, Robert Petryszak, Carmen Sancho-Serra, Agnes Oszlanczi, Cecilia Icoresi Mazzeo, Sophie Hoffman, Michał Krassowski, Kurtis Garbutt, Iva Kelava, Kezia Gaitskell, Slaveya Yancheva, Ee Von Woon, Victoria Male, Ingrid Granne, Karin Hellner, Krishnaa T Mahbubani, Kourosh Saeb-Parsy, Mohammad Lotfollahi, Elena Prigmore, Jennifer Southcombe, Rebecca A Dragovic, Christian M Becker, Krina T Zondervan, Roser Vento-Tormo

**Affiliations:** Wellcome Sanger Institute, Cambridge, UK; Oxford Endometriosis Care Centre, Nuffield Department of Women’s & Reproductive Health, University of Oxford, Oxford, UK; Wellcome Centre for Human Genetics, University of Oxford, Oxford, UK; Nuffield Division of Clinical Laboratory Sciences, Radcliffe Department of Medicine, University of Oxford, Oxford, UK; Department of Cellular Pathology, John Radcliffe Hospital, Oxford, UK; Department of Metabolism, Digestion and Reproduction, Institute of Developmental Reproductive and Developmental Biology, Imperial College London, London, UK; The Fertility Centre, Chelsea and Westminster Hospital, London, UK; Department of Haematology, University of Cambridge, Cambridge, UK; Cambridge Biorepository for Translational Medicine (CBTM), NIHR Cambridge Biomedical Research Centre, Cambridge, UK; Department of Surgery, University of Cambridge, Cambridge, UK

## Abstract

The human endometrium, the inner lining of the uterus, exhibits complex, dynamic changes throughout the menstrual cycle in response to ovarian hormones. Aberrant response of endometrial cells to hormones is associated with multiple disorders, including endometriosis. Previous single-cell studies of the endometrium profiled a limited number of donors and lacked consensus in defining cell types and states. Here, we introduce the Human Endometrial Cell Atlas (HECA), a high-resolution single-cell reference atlas, combining published and newly generated single-cell transcriptomics datasets of endometrial biopsies of women with and without endometriosis. The HECA assigned consensus cell types and states, and uncovered novel ones, which we mapped in situ using spatial transcriptomics. We quantified how coordinated interactions between cell states in space and time contribute to endometrial regeneration and differentiation. In the continuously changing *functionalis* layer, we identified an intricate coordination of TGFβ signalling between stromal and epithelial cells, likely crucial for cell differentiation. In the *basalis* layer, we defined signalling between fibroblasts and a new epithelial cell population expressing epithelial stem/progenitor markers, suggesting their role in endometrial regeneration. Additionally, integrating the HECA single-cell data with genome-wide association study data and comparing endometrial samples from women with and without endometriosis, we pinpointed subsets of decidualised stromal cells and macrophages as the most dysregulated cell states in endometriosis. Overall, the HECA is an invaluable resource for studying endometrial physiology, investigating endometrial disorders, and guiding the creation of endometrial microphysiological *in vitro* systems.

## Main

Human reproduction depends on the endometrium, the inner mucosal lining of the uterus. It prepares an optimal environment for embryo implantation and supports pregnancy if implantation is successful. In the absence of a pregnancy, the endometrium sheds each month during menstruation. Morphologically, the endometrium is composed of two layers: the ever-changing *functionalis* (adjacent to the uterine cavity) and the relatively constant *basalis* (adjacent to the myometrium). In response to ovarian steroid hormones, the *functionalis* undergoes repeated cycles of shedding and repair without scarring, extensive growth and differentiation^1,2^.

At the cellular level, the endometrium has a heterogeneous architecture. The endometrial epithelium consists of a horizontally interconnected network of *basalis* glands ^3–5^ contiguous with coiled *functionalis* glands extending vertically towards the uterine cavity, where a layer of *functionalis* luminal cells lines the endometrial surface. The *basalis* glands harbour epithelial stem/progenitor cells needed to regenerate the *functionalis* layer after menstruation^6–10^. The *functionalis* epithelium provides a site for embryo implantation, and produces secretions to nourish it. Stromal, fibroblast, perivascular and endothelial cells provide support and structural integrity, including rich vasculature within the tissue. An array of immune cells play crucial roles in endometrial shedding and repair^11,12^, as well as embryo implantation^13^. Cell-cell communication between the endometrial cells is key in maintaining tissue homeostasis and menstrual cycle progression.

During female reproductive years, the endometrium is highly heterogeneous, both inter- and intra-individually, and thus a large sample size is required to account for the dynamic changes it undergoes both in time (across the menstrual cycle) and space (across different tissue microenvironments). In recent years, several foundational studies atlasing the cellular composition of the human endometrium in health and pathologies with single cell^14–21^ and spatial^15–17^ technologies have been published. However, these cell censuses have so far profiled a limited number of samples, lacked even coverage of the menstrual cycle phases, and lacked consensus cell state annotation and reproducible marker gene signatures. In addition, they varied considerably in terms of clinical and phenotypic characterisation of the individuals from whom the samples were obtained. These factors have complicated comparisons across studies, with, for example, inconsistencies in the identification and naming of epithelial and stromal cell states. An integrated single-cell reference atlas of the human endometrium, encompassing the widest possible range of cell states and samples, is now warranted.

Endometrial heterogeneity is further increased by endometrial/uterine disorders which are highly prevalent globally. For example, abnormal menstrual bleeding affects up to a third of all women in their lives, ~417,000 new cases of endometrial cancer are diagnosed yearly, and ~190 million women world-wide suffer from endometriosis^22–24^. In endometriosis, endometrial-like cells grow outside of the uterus (i.e. ectopically), and are associated with debilitating chronic pain and subfertility that can have a substantial negative impact on quality of life^25^. Conflicting evidence exists about whether and to what extent the endometrium itself (i.e. the eutopic endometrium) differs between those with and without endometriosis^26,27^. Recently, single-cell studies analysing small sample sizes, reported dysregulation of the stromal and immune compartments in the endometrium of women with endometriosis to various degrees^16,18,20,28,29^. Larger sample sets are now needed if we are to unpick whether and how the endometrium differs in those with and without the condition. In this context, well-annotated reference cell atlases can provide invaluable insights.

Here, we assemble a consensus cell atlas of the endometrium, the Human Endometrial Cell Atlas (HECA), by harmonising the transcriptomic and donor metadata information of ~626,000 cells and nuclei from previously published and newly generated datasets (https://www.reproductivecellatlas.org/). We identify new cell populations, including an epithelial *CDH2+* population in the *basalis* and distinct populations of *functionalis* epithelial and stromal cells characteristic of the early secretory phase. We describe the molecular signals likely mediating the spatiotemporal organisation and function of cellular niches throughout the menstrual cycle and provide a new interactive portal to visualise and query the predicted cell-cell communication. Finally, we use the HECA to give cellular context to genetic associations identified by the largest endometriosis genome-wide association study (GWAS) meta-analysis^30^. This analysis identifies macrophages and subsets of decidualised stromal cells as the endometrial cell types expressing the genes affected by the variants associated with endometriosis.

## Results

### Harmonised data to generate the HECA

To comprehensively define endometrial cell types and states and how they change across the menstrual cycle, we analysed a total of ~626,000 high-quality cells and nuclei from 121 individuals (**Figure 1a-b**). We started by creating a single-cell reference atlas, which we termed the HECA (**Figure 1c**). To create the HECA, we integrated six publicly available single-cell RNA sequencing (scRNA-seq) datasets (Wang et al.^14^, Garcia-Alonso et al.^15^, Tan et al.^16^, Lai et al.^19^, Fonseca et al.^17^, Huang et al.^18^) with our newly generated dataset (termed Mareckova (cells) dataset) (**Figure 1b**). Harmonisation of metadata across the studies and application of strict data quality control filters (**see Methods**) was essential for the integration. The final integrated HECA consisted of ~314,000 high-quality cells from 7 datasets, of which ~76,000 cells were newly profiled by us (**Supplementary Table 2**). It included a total of 63 individuals both with endometriosis (i.e cases) and without endometriosis (i.e. controls), with samples collected either during natural cycles or when taking exogenous hormones (**Figure 1b & c, Supplementary Table 1**). The majority of samples analysed were superficial biopsies of the endometrium, predominantly sampling the *functionalis* layer from living donors. Three samples from the uteri of donors who died of non-gynaecological causes contained full-thickness endometrium, encompassing both the *functionalis* and *basalis* layers, with attached subjacent myometrium.

**Figure 1.**
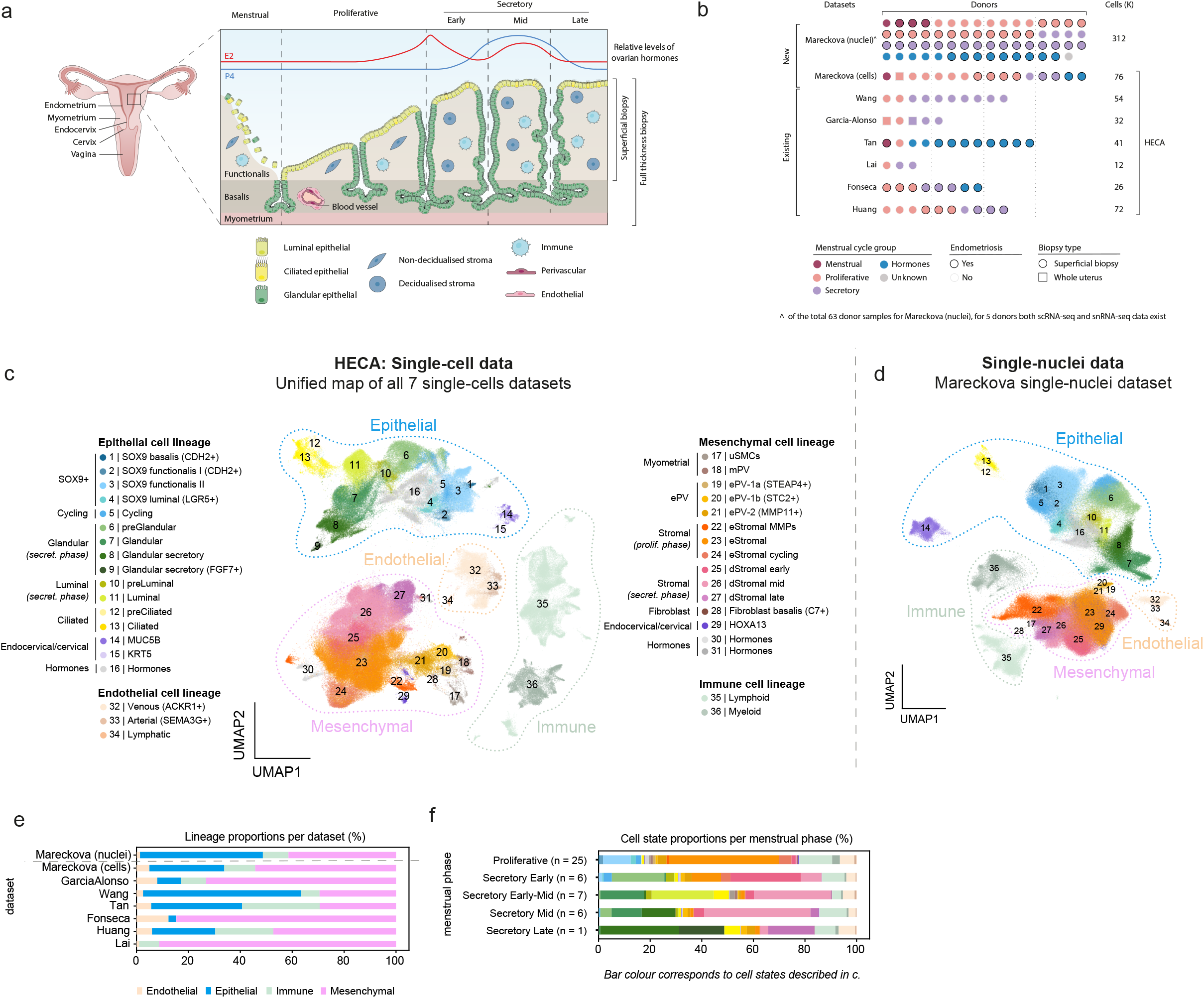
Harmonised cellular map of the human endometrium. **a,** Schematic illustration of the human uterus and cellular composition of the endometrium as it undergoes morphological changes across the menstrual cycle. **b**, List of datasets analysed and contribution of the number of donors, cells/nuclei, endometrial histology and endometriosis status of all samples profiled per dataset. **c,** UMAP projections of scRNA-seq data from a total of 63 individuals and ~314,000 cells coloured by cell state. **d,** UMAP projections of snRNA-seq data from a total of 63 individuals and ~312,000 nuclei coloured by cell state. **e,** Bar plot showing the contribution of each of the scRNA-seq datasets to the main cellular lineages (endothelial, epithelial, immune and mesenchymal lineages) as shown in c. **f,** Bar plot showing the cellular composition of endometrial biopsies from the proliferative (*n* = 25), early secretory (*n* = 6), early/mid secretory (*n* = 7), mid secretory (*n* = 6) and late secretory (*n* = 1) phases of the menstrual cycle for the scRNA-seq data presented in c. dStromal, decidualised stromal cells; eStromal, endometrial stromal cells specific to proliferative phase; HECA, human endometrial cell atlas; MMPs, matrix metalloproteinases; mPV, myometrial perivascular cells; ePV, endometrial perivascular cells; scRNA-seq, single-cell RNA-sequencing; secret., secretory; snRNA-seq, single-nucleus RNA-sequencing; UMAP, uniform manifold approximation and projection; uSMCs, uterine smooth muscle cells.

We observed striking differences between the cellular composition of the integrated scRNA-seq datasets, with variable recovery of epithelial, mesenchymal, endothelial and immune cells (Figure 1e). Choice of tissue digestion protocol, sampling bias (technical variation), menstrual cycle stage and use of exogenous hormones (biological variation) could all be responsible for the differences observed (**see Methods & Supplementary Figure 1 & Supplementary Table 1**). The dataset-specific cellular proportions prompted us to generate an independent single-nucleus RNA sequencing (snRNA-seq) dataset for 63 additional donors (**Figure 1b & d**). The large number of individuals in this dataset allowed us to overcome the technical variation introduced when data are generated by different laboratories. We profiled ~312,000 high-quality nuclei from snap-frozen samples of superficial endometrial biopsies (**Figure 1b & d, Supplementary Figure 2, Supplementary Table 2**), collected during natural cycles, when taking exogenous hormones, and included samples for donors with and without endometriosis (**Figure 1b**). Together, this dataset represents the largest set of human endometrial samples profiled at the single-cell/-nucleus transcriptomic level by a single laboratory so far. To align the cell state annotations across the scRNA-seq and snRNA-seq datasets, and determine the robustness of the HECA, we transferred cell states labels between datasets using machine learning tools (**see Methods**). Out of the endometrial cells identified by scRNA-seq, the majority were validated in the nuclei dataset (**Supplementary Figure 2b-c**).

As expected, the majority of the cell populations were of endometrial origin, but the atlas also contained populations exclusively present in the myometrium from the whole uterine samples (e.g. uterine smooth muscle cells (uSMCs) and myometrial perivascular cells (mPV)). In addition, we detected a small number of mesenchymal HOXA13+ and epithelial KRT5+ cells, which based on their marker gene expression were likely cervical cell contamination. This was supported by their transcriptomic similarity to cervical cells when we compared the HECA with a publicly available scRNA-seq dataset of the cervix^31^(**Supplementary Figure 1e-i**). We did not detect any endometriosis-specific cell state in neither the scRNA-seq nor snRNA-seq data, providing further evidence that at the cellular level of the endometrium, differences between controls and cases may be more subtle. However, additional cell states appeared in samples from donors taking exogenous hormones, indicating that exogenous hormones strongly impact the global transcriptome of epithelial cells, an observation supported by both data sources (**Supplementary Figure 3**).

Altogether, we generated the most comprehensive reference atlas of the human endometrium (i.e. the HECA), which can now be used to map and contextualise newly processed samples and external datasets using the transfer learning framework scArches^32^. To facilitate this process, we prepared computational tutorials (**see Methods**) and provide the weights from the trained scANVI model^33^ of the HECA available at https://www.reproductivecellatlas.org/.

### Spatiotemporal complexity of the endometrial epithelium

The endometrial epithelium consists of a complex network of *basalis* glands, which house the stem/progenitor cells needed to regenerate the *functionalis* glands extending into the uterine cavity, lined by a layer of luminal cells (**Figure 1a**). Here, we characterised, with fine granularity, the cell states forming the different regions of the endometrial epithelium across the proliferative and secretory phases of the menstrual cycle.

We identified a novel population, the SOX9 basalis (*CDH2*+) cells, that was not reported by previous single-cell transcriptomics atlases. These cells expressed markers described for endometrial epithelial stem/progenitor cells (*SOX9*, *CDH2, AXIN2, ALDH1A1*^9,34,35^)(**Figure 2a**). Using spatial transcriptomics and single molecule fluorescence in situ hybridisation (smFISH) imaging, we mapped this population to the *basalis* glands region in full thickness endometrial biopsies from both proliferative and secretory phases (**Figure 2b-c**). Cell-cell interaction analyses indicated that the SOX9 basalis (*CDH2+*) population interacts with the fibroblast basalis (i.e. Fibroblast basalis *C7*+) population via the expression of *CXCR4* and *CXCL12*, respectively (**Figure 2d**). The *CXCL12/CXCR4* axis is known to have a role in the maintenance of the stem cell niche in other tissues^36^, providing further evidence for the stem/progenitor nature of this cell subset.

**Figure 2.**
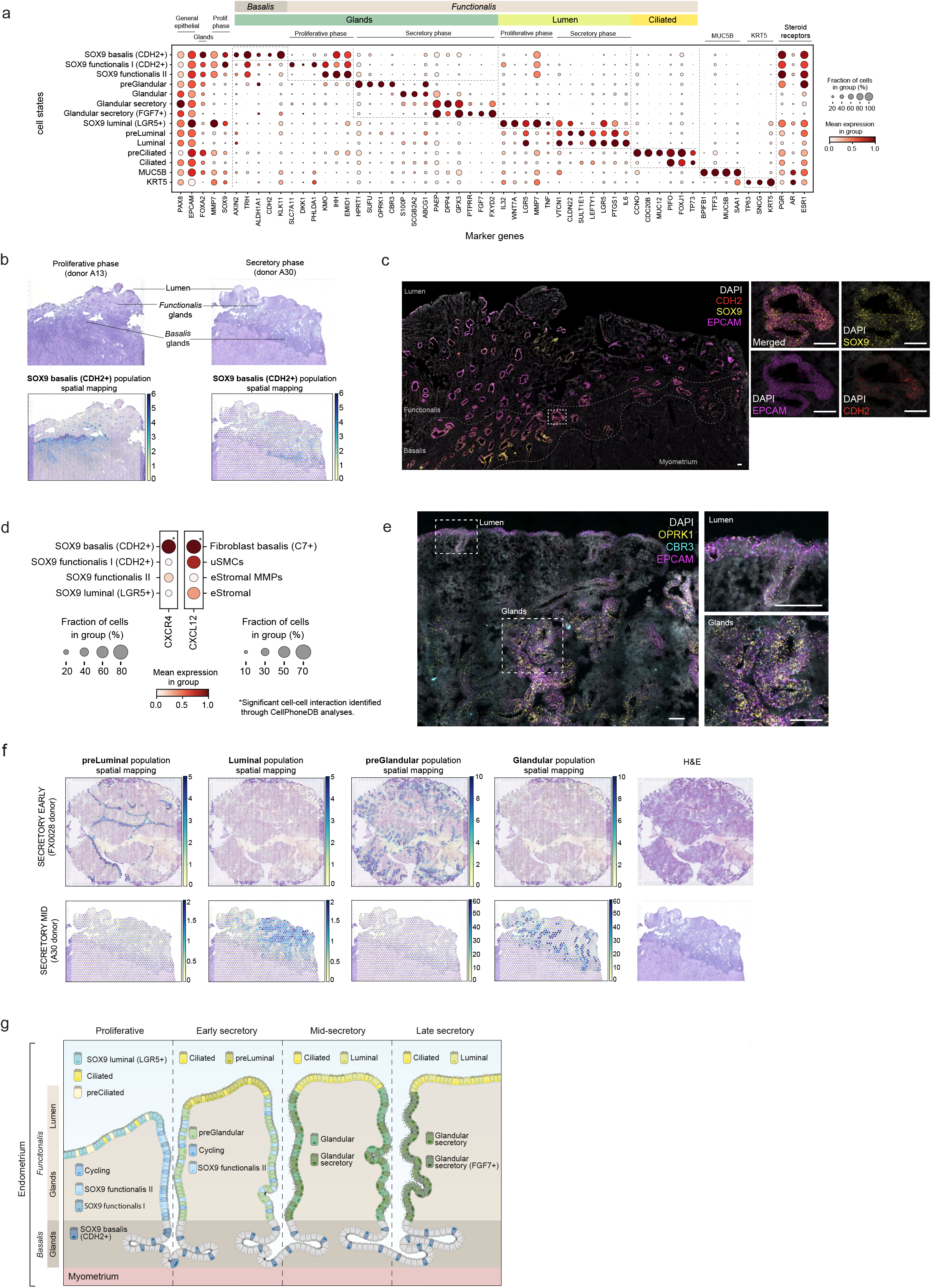
Spatiotemporal complexity of epithelial cells. **a,** Dot plot showing normalised, log-transformed and variance-scaled expression of genes (*x*-axis) characteristic of the identified epithelial cell states (*y*-axis) in scRNA-seq data. **b**, Visium spatial transcriptomics data and an H&E image of the same tissue section are shown. Spot colour indicates estimated cell state density for the SOX9 basalis (CDH2+) population in each Visium spot, as computed by cell2location. Spatial mapping of the SOX9 basalis (CDH2+) population is shown in sections of whole-uterus biopsies from donors A13 (proliferative phase) and A30 (secretory phase). **c,** High-resolution multiplexed smFISH of a section of a whole-uterus biopsy from donor A13 stained for *DAPI* (white, nuclei), *EPCAM* (magenta, epithelial cells), *SOX9* (yellow, epithelial cells), *CDH2* (red, epithelial cells). The dotted line highlights the *basalis* area of the endometrium where signal for all markers and their co-localisation is detected within the epithelial glands. The inset shows a representative zoom-in of one of the glands and signal co-localisation. Scale bars = 100 µm. **d,** Dot plot showing normalised, log-transformed and variance-scaled expression of *CXCR4* and *CXCL12* (*x*-axis) in a selection of epithelial and mesenchymal cells (*y*-axis) in scRNA-seq data. Asterisk denotes a significant cell-cell interaction identified through CellPhoneDB analyses. **e,** Left, high-resolution multiplexed smFISH of a section of a superficial biopsy from donor FX1233 showing the expression of *DAPI* (white, nuclei), *EPCAM* (magenta, epithelial cells), *CBR3* (cyan, preGlandular cells), and *OPRK1* (yellow, preGlandular cells). The dashed outlines indicate areas shown magnified to the right. Top right, a magnified image of the luminal region with low *OPRK1* and *CBR3* signal. Bottom right, a magnified image of the glandular region with high and co-localised *OPRK1* and *CBR3* signal. Scale bars = 100 µm. **f**, Visium spatial transcriptomics data and an H&E image of the same tissue section are shown. Spot colour indicates estimated cell state density for the preLuminal, Luminal, preGlandular and Glandular populations in each Visium spot, as computed by cell2location. Spatial mapping of preLuminal, Luminal, preGlandular and Glandular populations is visualised in a section of a superficial biopsy from donor FX0028 (early secretory phase) and a section of a whole-uterus biopsy from donor A30 (mid secretory phase). **g,** Schematic illustration of the spatiotemporal complexity of the endometrial epithelium across the proliferative and secretory phases. eStromal, endometrial stromal cells specific to proliferative phase; MMPs, matrix metalloproteinases; Prolif., proliferative; smFISH, single molecule fluorescence in situ hybridisation; uSMCs, uterine smooth muscle cells.

The cellular composition of the *functionalis* glands showed highly dynamic changes across the proliferative and secretory phases (**Figure 2a**). During the proliferative phase, we uncovered further heterogeneity within the known SOX9+ cell population^15^. Specifically, we identified two SOX9+ subpopulations: SOX9 functionalis I and II, which we mapped to the functionalis glands (**Supplementary Figure 4a**). The SOX9 functionalis I population expressed *CDH2*, high levels of *SOX9* and was marked by the expression of *PHLDA1* and *SLC7A11.* The SOX9 functionalis II population exhibited lower expression of *SOX9*, was negative for *CDH2* and distinctly expressed *KMO*, *IHH* and *EMID1*. The luminal proliferative epithelium was defined by the presence of SOX9 luminal (*LGR5*+), pre-ciliated and ciliated cells (**Figure 1f**, **2a**), as previously described by us^15^. As expected, we also detected a larger proportion of cycling epithelial cells in the proliferative phase endometrium (**Figure 1f**).

During the secretory phase, the SOX9+ populations were markedly reduced as the endometrium underwent further differentiation in order to prepare a receptive environment for blastocyst implantation (**Figure 1f**). During the differentiation process, both the glandular and luminal epithelium undergo dramatic transcriptomic and morphological changes. Having a larger number of samples allowed us to further subdivide the secretory phase into early, early-mid, mid and late secretory phases and define the populations associated with these stages (**Figure 1f, see Methods**). For the first time, we uncovered the transcriptomic profiles of cells characteristic of the *functionalis* layer during the early secretory phase (i.e. the preGlandular and preLuminal populations; **Figure 2a & e-f**). These populations were transcriptomically similar to the previously described glandular and luminal populations^15^, but appeared at earlier stages of the cycle and expressed markers not defined previously. For the preGlandular population these included *OPRK1, SUFU, CBR3, HPRT1*, and for the preLuminal population *SULT1E1* was the most specific marker (**Figure 2a**). Using spatial transcriptomics, we confidently mapped both populations to early, but not mid-secretory samples. Specifically, the preLuminal population mapped to the lumen and the preGlandular population to the *functionalis* glands (**Figure 2f & Supplementary Figure 4b**). We further confirmed the preGlandular subset using smFISH imaging (**Figure 2e & Supplementary Figure 4c**).

The number of preGlandular and preLuminal cells decreased in the early-mid and mid-secretory phase samples, with the dominant cell states being the previously described glandular, luminal and ciliated populations^15^(**Figure 1f**). Lastly, analysing a single sample profiled from the late secretory phase, we observed the presence of a glandular secretory population that upregulated *FGF7*, a mitogen found to have a role in wound healing in other contexts^37,38^.

We detected a previously described population of MUC5B epithelial cells^16^ expressing *MUC5B, TFF3, SAA1, BPIFB1*. As in previous studies^16^, we also observed varied expression of the cell type marker *MUC5B* when staining full-thickness endometrial biopsies using smFISH (**Supplementary Figure 4d**). However, when projecting a publicly available scRNA-seq dataset of the cervix^31^ onto our HECA (**Supplementary Figure 1h**), we found a cluster of cervical epithelial cells matching the transcriptome of this population (**Supplementary Figure 1g-i**). This result implies the *MUC5B+* cells are likely to be present in the endocervical columnar epithelial cells^31,39^, and we cannot disregard the possibility that in the HECA, the MUC5B population comes exclusively from the endocervix.

In summary, we defined and spatially located novel epithelial cell states across the proliferative and secretory phases, including a putative stem/progenitor cell population found within the *basalis* layer and multiple transitory cell states dominating the *functionalis* layer.

### Stromal-epithelial crosstalk across the menstrual cycle

During the menstrual cycle, stromal and epithelial cells synchronise their differentiation under the influence of ovarian hormones, as well as locally produced paracrine factors. Here we used the HECA’s fine-grained classification of stromal and epithelial cell states across the menstrual cycle to infer cell-cell communication occurring *in vivo* along the endometrial cellular niches in space (i.e. *basalis*, *functionalis*) and time (i.e. menstrual cycle phase).

Within the *functionalis* layer, endometrial stromal cells (eStromal) specific to the proliferative phase and decidualised stromal cells (dStromal) specific to the secretory phase were defined previously at the single-cell level^15,40^. In the HECA, we further identified a new type of eStromal cells (eStromal MMPs) in samples collected during the menstrual and early proliferative phases (**Figure 3a, Supplementary Figure 1d**), characterised by the upregulation of metalloproteases (*MMP1, MMP10, MMP3*) and inhibin A (*INHBA*) (**Figure 3a**).

**Figure 3.**
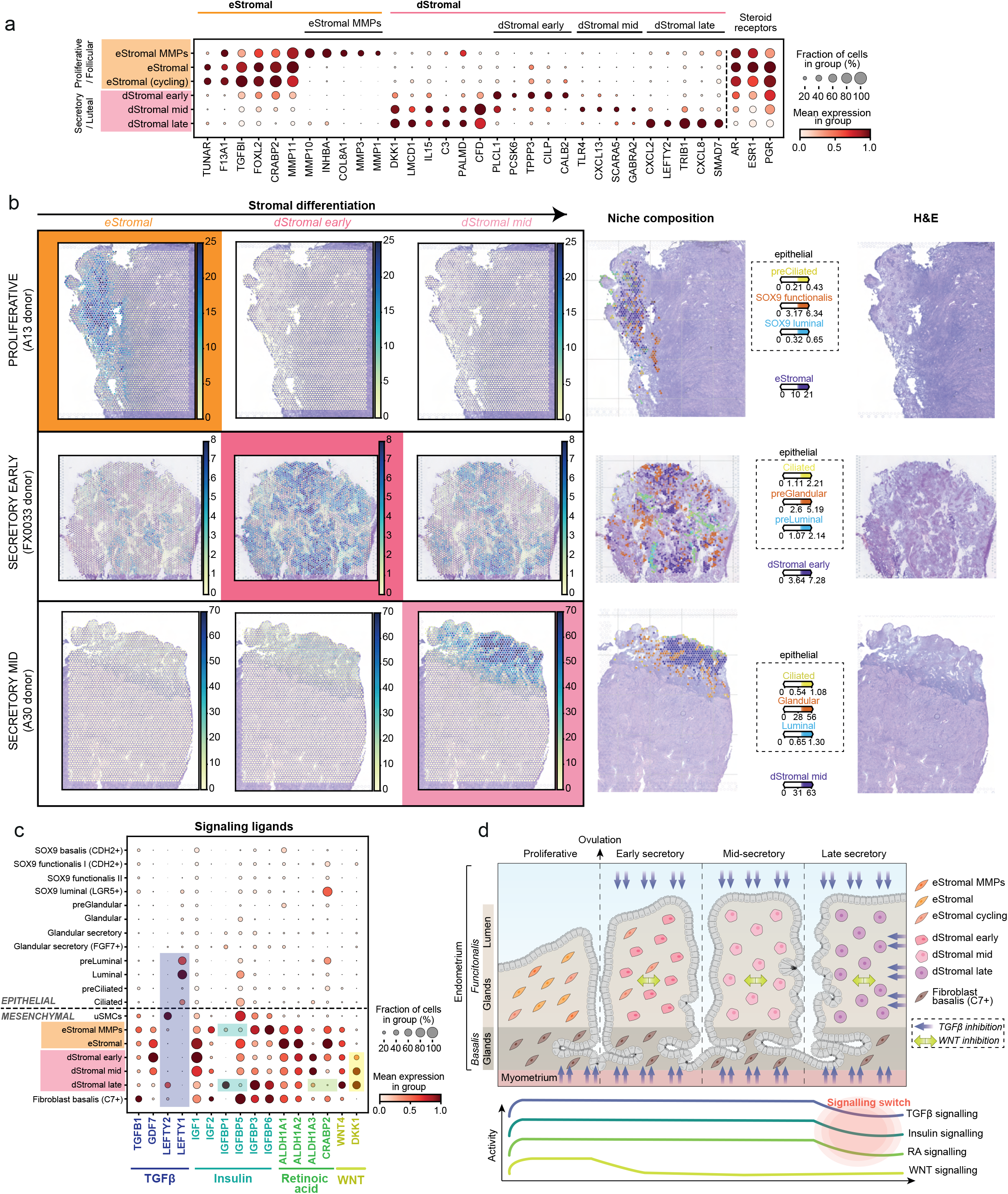
Endometrial stromal cell heterogeneity and stromal-epithelial cell cross-talk across the menstrual cycle. **a,** Dot plot showing normalised, log-transformed and variance-scaled expression of genes (*x*-axis) characteristic of the identified stromal cell states (*y*-axis) in scRNA-seq data. **b,** Visium spatial transcriptomics data and an H&E image of the same tissue section are shown. Spot colour indicates estimated cell state density for a specific cell population in each Visium spot as computed by cell2location. Spatial mapping of the eStromal, dStromal early and dStromal mid cell populations is shown in a section of a whole-uterus biopsy from donor A13 (top panel, proliferative phase), a section of a superficial biopsy from donor FX0033 (middle panel, early secretory phase) and a section of a whole-uterus biopsy from donor A30 (bottom panel, mid secretory phase). Mapping of menstrual cycle phase-relevant epithelial cell populations is also shown in the niche composition panel. **c,** Dot plot showing normalised, log-transformed and variance-scaled expression of genes (*x*-axis) in epithelial and mesenchymal cell states (*y*-axis) in scRNA-seq data. Signalling ligands involved in TGFβ, insulin, retinoic acid and WNT signalling are shown. **d,** Schematic illustration of the temporal complexity of endometrial stromal cells and signalling pathways across the proliferative and secretory phases. eStromal, endometrial stromal cells specific to proliferative phase; dStromal, decidualised stromal cells; MMPs, matrix metalloproteinases; RA, retinoic acid; TGFβ, transforming growth factor beta.

In secretory phase samples, we identified three new dStromal cell states appearing at different stages of the secretory phase. Early decidualised stromal cells (dStromal early) were enriched in the early secretory phase samples and upregulated the progesterone-induced gene *PLCL1*^41^ (**Figure 3a-b**). The mid decidualised stromal population (dStromal mid) mapped to early-mid and mid-secretory phase samples and upregulated *DKK1* (**Figure 3a-b**), a WNT-inhibitor crucial for the differentiation of epithelial secretory glands^15^. Late decidualised stromal cells (dStromal late) were present in both mid- and late secretory phase samples (**Figure 1f**) and upregulated the premenstrual marker *LEFTY2*^42^ (**Figure 3a**). Both the dStromal mid and late populations downregulated oestrogen and progesterone receptors (*ESR1* and *PGR*).

We uncovered an intricate spatiotemporal regulation of transforming growth factor beta (TGFβ) signalling (**Figure 3c**). Specifically, the TGFβ superfamily receptors were ubiquitously expressed by all epithelial and stromal cells at all stages of the menstrual cycle (**Supplementary Figure 5**). Meanwhile, the ligands of TGFβ and growth differentiation factor (GDF) subfamilies (*TGFB1* and *GDF7*, respectively) were upregulated by all stromal cells until mid/late secretory phase, when expression dropped (**Figure 3c**). Interestingly, the activity of TGFβ signalling appeared confined within specific spatial and temporal boundaries by its antagonists, *LEFTY1* and *LEFTY2*. On one hand, *LEFTY1* was expressed by epithelial cells of the lumen (ciliated and luminal) and *LEFTY2* by uSMCs of the myometrium (**Figure 3c**). This pattern of expression likely establishes a top-bottom spatial boundary of TGFβ activity. On the other hand, the temporal boundary seemed to be determined by the expression of *LEFTY2* as well as *SMAD7* (the inhibitor of SMADs, downstream effectors of TGFβ). These two molecules were expressed by the dStromal late population (**Figure 3a**), suggesting TGFβ activity is switched off towards the end of the menstrual cycle (**Figure 3d**). Additionally, using our detailed cell annotation, we could pinpoint the specific stromal cell states involved in previously reported stromal-to-epithelial cell signalling mediated by WNT inhibition^40^, insulin^43^ and retinoic acid^44^ across the menstrual cycle (**Figure 3c & d**).

Taken together, our data supported a rise in TGFβ, insulin, WNT and retinoic acid signalling from early stages of the proliferative phase (**Figure 3d**). WNT inhibition marked the beginning of the secretory phase with the initiation of stromal cell decidualisation. In the late secretory phase, our data supported a signalling switch in the use of TGFβ signalling, insulin growth factors and retinoic acid metabolism (**Figure 3c & d**). The full collection of cell-cell communication factors, identified through CellPhoneDB analyses^45^ can be visualised and queried using our new interactive portal at https://www.reproductivecellatlas.org.

### Macrophages in endometrial regeneration

To gain insights into the diversity and dynamics of innate immune cells in the endometrium and pinpoint their involvement in the regeneration process, we examined our comprehensive datasets (*n* = 32,322 cells and *n* = 24,820 nuclei). These datasets captured the three uterine Natural Killer cell populations (uNK1, uNK2, uNK3) and the two uterine macrophage populations (uM1 and uM2) previously identified by us in the endometrium during pregnancy (i.e. decidua)^46^ (**Figure 4a, Supplementary Figure 6a-e**). Differential cell abundance analysis (**see Methods**) demonstrated an increase in the abundance of uNK1 cells during the secretory phase, in line with previous reports of granular endometrial immune cells proliferating during the secretory phase^47,48^ (**Figure 4b, Supplementary Figure 6f**). Cell abundance of the other immune cell types did not differ between the proliferative and secretory phases.

**Figure 4.**
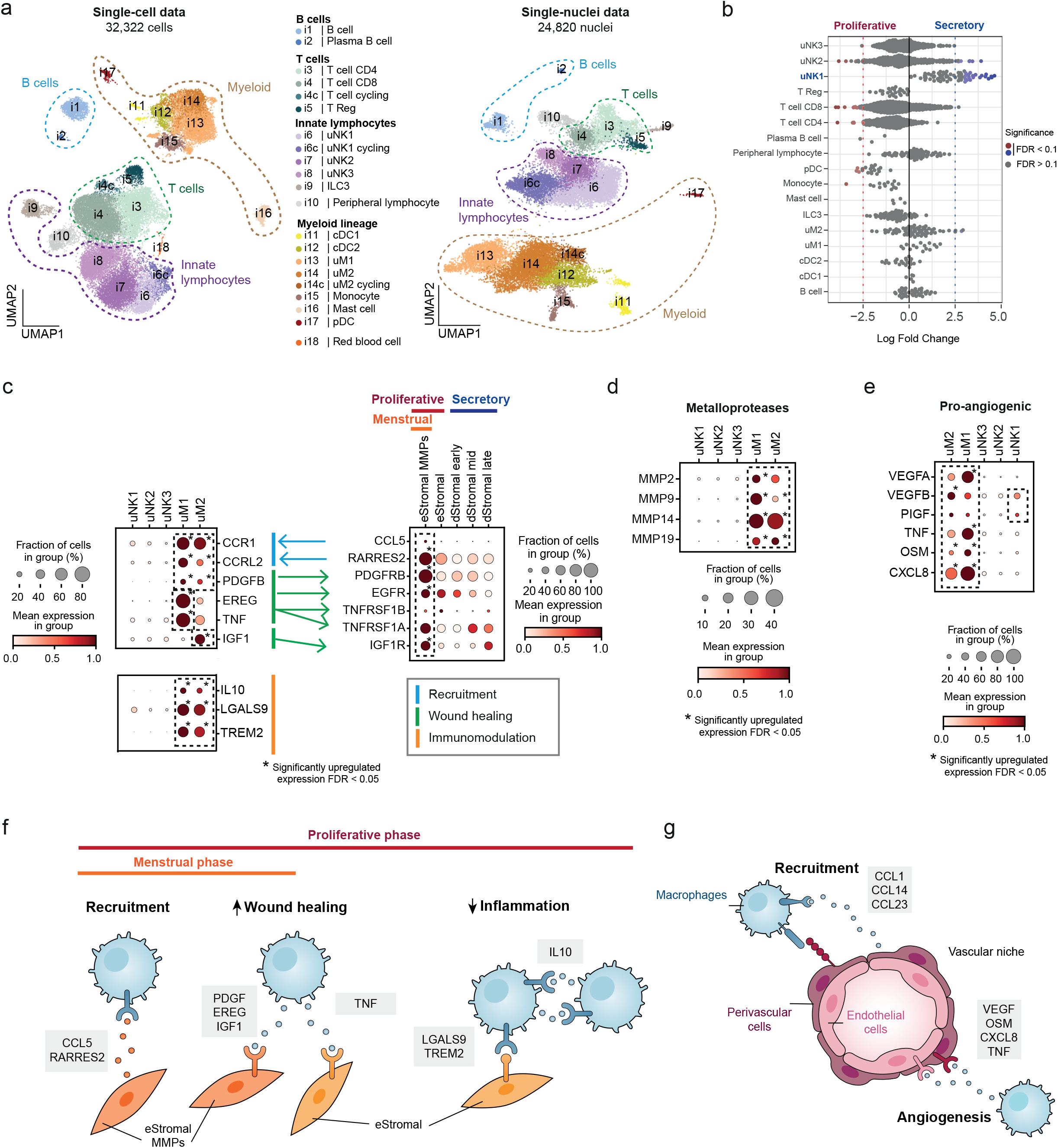
Predicted ligand-receptor interactions and role of macrophages in endometrial repair and regeneration. **a,** Left, UMAP projections of scRNA-seq data for 32,322 immune cells coloured by cell type. Right, UMAP projections of snRNA-seq data for 24,820 immune cells/nuclei coloured by cell type. **b,** Beeswarm plot of the distribution of log fold change across the menstrual cycle (proliferative and secretory phases) in neighbourhoods containing immune cells from different cell type clusters in scRNA-seq data. Differentially abundant neighbourhoods at log fold change > 2.5 and spatial FDR < 0.1 are coloured. **c,** Dot plot showing normalised, log-transformed and variance-scaled expression of genes (*y*-axis) in uNK and uM cell states (*x*-axis) in scRNA-seq data. Asterisk denotes significantly upregulated expression at FDR < 0.05. **d,** Dot plots showing normalised, log-transformed and variance-scaled expression of signalling molecules and receptors (*y*-axes) upregulated in uNK, uM and stromal cell states (*x*-axes) in scRNA-seq data. Asterisk denotes significantly upregulated expression at FDR < 0.05. The predicted cell-cell communication between uNK, uM and stromal cell states, including its likely role, is shown by differently coloured arrows. **e,** Dot plot showing normalised, log-transformed and variance-scaled expression of pro-angiogenic signalling molecules (*y*-axis) upregulated in uNK and uM cell states (*x*-axis) in scRNA-seq data. Asterisk denotes significantly upregulated expression at FDR < 0.05. **f,** Schematic illustration of macrophage and stromal cell signalling during the menstrual and proliferative phases, likely involved in macrophage cell recruitment, increasing wound healing abilities and dampening inflammation in stromal cells. **g,** Schematic illustration of macrophage, endothelial cell and perivascular cell signalling likely involved in macrophage recruitment and angiogenesis. Cells from donors on hormones and donors with endometriosis were excluded from analyses shown in **b - e** of this figure. cDC, conventional dendritic cells; dStromal, decidualised stromal cells; eStromal, endometrial stromal cells specific to proliferative phase; FDR, false discovery rate; ILC3, innate lymphoid cell type 3; MMPs, matrix metalloproteinases; pDC, plasmacytoid dendritic cell; scRNA-seq, single-cell RNA-sequencing; snRNA-seq, single-nucleus RNA-sequencing; T Reg, T regulatory cells; UMAP, uniform manifold approximation and projection; uM, uterine macrophages; uNK, uterine natural killer cells.

To deepen our understanding of the role innate immune cells play in endometrial regeneration, we interrogated their cell-cell communication with stromal, endothelial and perivascular (PV) cells. We focused on significantly upregulated genes in uMs and uNK cells when compared to all immune cell subsets, especially during the menstrual and proliferative phase, a period when these innate immune cells are thought to aid in endometrial wound healing and regrowth (**see Methods**). We found that the eStromal MMPs population (characteristic of the menstrual phase) expressed integrins and cytokines (*CCL5, RARRES2)* which can bind their cognate receptors upregulated by uMs (*CCR1, CCRL2*) (**Figure 4c; Supplementary Figure 7a**). This interaction likely supports the previously described recruitment of uMs to the tissue during menstruation^49,50^. We also noticed that uMs upregulated *PDGFB,* a protein from the PDGF family, known for their role in wound healing and repair in various tissues^51,52^. In the endometrium, it could operate by binding to the *PDGFRB* receptor, which is upregulated by eStromal MMPs (**Figure 4c**). Additionally, uMs upregulated *TNF* (uM1), as well as growth factors such as *IGF1* (uM2) and *EREG* (uM1). These could stimulate the proliferation and survival of eStromal MMPs and proliferative eStromal cells by binding to their corresponding receptors (*EGFR, TNFRSF1A, TNFRSF1B* and *IGF1R)* (**Figure 4c**). Finally, uMs also expressed immunoregulatory genes (*IL10, LGALS9, TREM2*) that could enhance anti-inflammatory responses in the proliferative phase endometrium required for the characteristic scarless regeneration of this tissue (**Figure 4c**).

Additionally, angiogenesis is also critical for tissue repair, and macrophages are known to play a role in this process^53^. In the endometrium, there is a profound growth of blood vessels during the proliferative phase as the *functionalis* regenerates and thickens after being shed. During the secretory phase, the vasculature further matures and coils in preparation for pregnancy. To investigate the potential interplay between uMs and the vasculature, we first defined the endometrial vascular niche. We identified three subsets of endothelial cells (venous, arterial and lymphatic) and three subsets of endometrial PV cells (ePV-1a expressing *STEAP4,* ePV-1b expressing *STC2*, and ePV-2 expressing *MMP11*) (**Supplementary Figure 6g-h**). ePV-2 exhibited transcriptomic similarities to endometrial stromal cells, suggesting a transitional population between PV and stromal cells (**Supplementary Figure 1c**).

Cell-cell communication analyses predicted signalling between the vasculature and uMs, and to a lesser extent also with uNK1 cells. Endothelial cells and ePV-1s expressed multiple extracellular matrix proteins (ECM) and cytokines (*CCL14, CCL23, CCL26*), which potentially could act to recruit innate immune cells (**Supplementary Figure 7b**). Additionally, PVs expressed *CSF1* (major macrophage growth factor), which could create a favourable environment for macrophages, stimulating their differentiation and function. In turn, uMs expressed multiple growth factor members of the pro-angiogenic VEGF family (*VEGFA, VEGFB, PIGF)*, and vascular remodelling factors (*TNF*^54^, *OSM*^55^, *CXCL8*^56^), whose cognate receptors (*NRP1, NRP2, FLT1, TNFRSF1A-B, OSMR, LIFR, ACKR1*) were expressed by the endothelial cells (**Figure 4e, Supplementary Figure 7b**). Among the innate lymphocytes, uNK1 was the only cell subset that expressed pro-angiogenic factors (*VEGFB* and *PIGF*), although at lower levels than uterine macrophages (**Figure 4e**).

Altogether, our analysis underscored macrophages as the major endometrial immune cells participating in the process of blood vessel formation, wound healing and anti-inflammatory responses (**Figure 4f-g**). The latter two processes are likely to aid the stromal cells in healing without scarring.

### Altered stromal-immune cell homeostasis in the eutopic endometrium of endometriosis cases

We next investigated whether cellular composition of the endometrium differs between endometriosis cases and controls during natural menstrual cycles, as we did not detect any endometriosis-specific cell types. After accounting for menstrual cycle phase (**see Methods**), differential abundance analysis of our nuclei dataset revealed lower abundance of decidualised stromal cells (dStromal mid) and higher abundance of uM1 macrophages in endometriosis cases (**Figure 5a**). Interestingly, decidualised stromal cells (dStromal early and dStromal mid) and macrophages (uM1 and uM2) were also identified as the top cell types enriched for the expression of genes positionally close to endometriosis risk variants when performing functional GWAS (fGWAS) analysis across the HECA cell types (**Figure 5b, see Methods**). The fGWAS analysis provided, for the first time, cellular context to a large-scale endometriosis GWAS meta-analysis^30^

**Figure 5.**
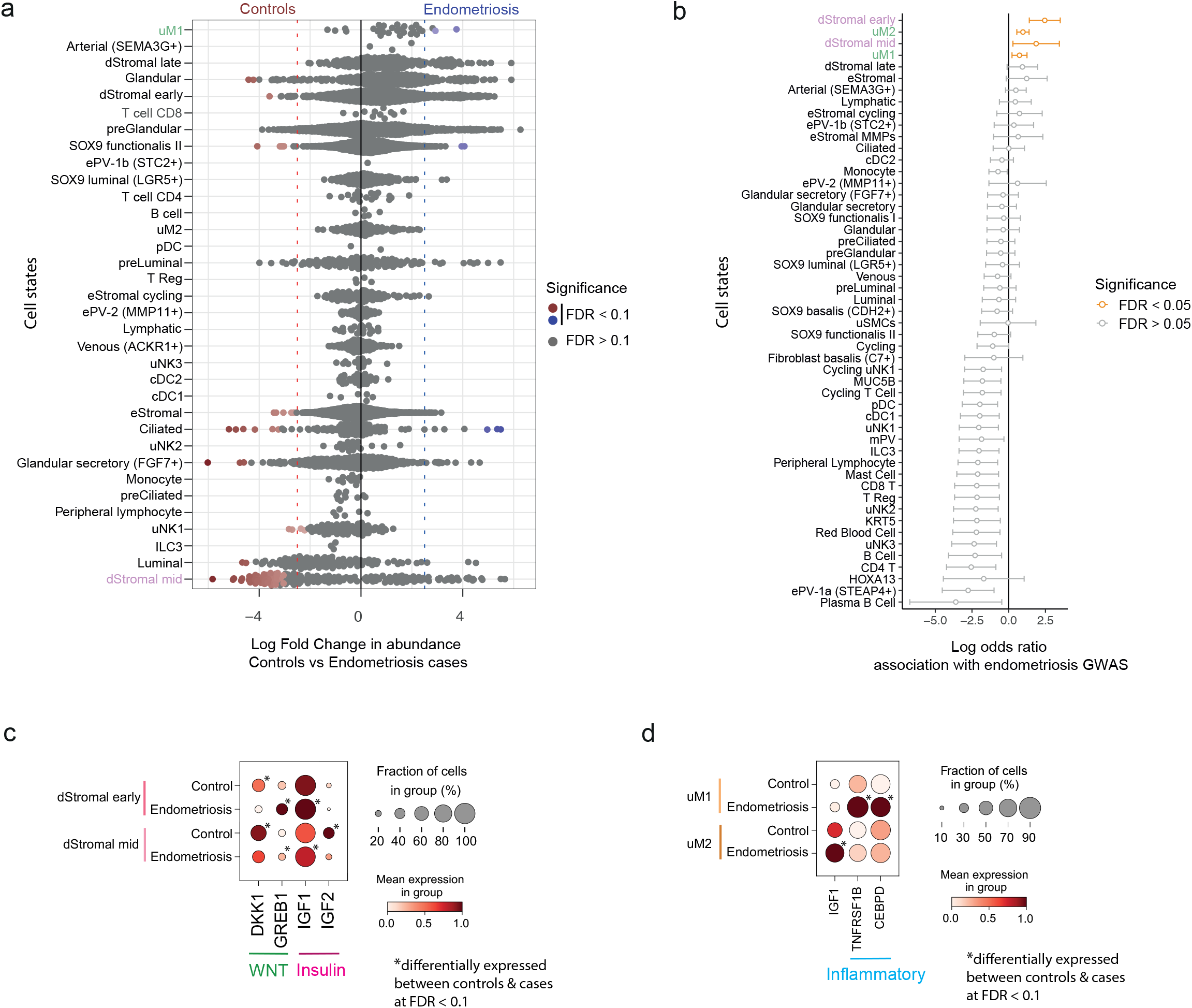
Endometrial stromal-immune cell niche in endometriosis. **a,** Beeswarm plot of the distribution of log fold change between conditions (controls and endometriosis cases) in neighbourhoods containing endometrial cells from different cell type clusters in snRNA-seq data. Differentially abundant neighbourhoods at log fold change > 2.5 and spatial FDR < 0.1 are coloured. **b,** Forest plot showing the log odds ratio (*x*-axis) of the enrichment for expression of genes associated with endometriosis in each endometrial cell type (*y*-axis). Cell types in orange have FDR < 0.05. **c,** Dot plot showing normalised, log-transformed and variance-scaled expression of differentially expressed genes (*x*-axis) in dStromal cell states of controls and endometriosis cases (*y*-axis) in scRNA-seq data. **d,** Dot plot showing normalised, log-transformed and variance-scaled expression of differentially expressed genes (*x*-axis) upregulated in uM cell states (*y*-axis) in scRNA-seq data. Cells from donors on hormones were excluded from all analyses shown in this figure. dStromal, decidualised stromal cells; FDR, false discovery rate; uM, uterine macrophages.

To further explore the four cell populations identified as endometriosis-relevant, we performed differential gene expression analyses between controls and endometriosis cases. In the stromal compartment of endometriosis cases, we observed changes in gene expression that are likely to alter the WNT and insulin signalling pathways (**Figure 5c**). Specifically, *GREB1* (a GWAS-linked gene induced by WNT signalling^57,58^) was significantly upregulated, while *DKK1* (WNT inhibitor) was significantly downregulated in both dStromal early and dStromal mid cells in endometriosis. These changes suggested sustained WNT signalling in the secretory phase endometrium of donors with endometriosis. Similarly, we observed a dysregulation of insulin growth factors *IGF1* (a GWAS-linked gene) and *IGF2.* In dStromal early and dStromal mid populations, *IGF1* was significantly upregulated, while *IGF2* was significantly downregulated in endometriosis cases. *IGF1* and *IGF2* play a role in cell proliferation and differentiation^59,60^, suggesting dysregulation of these processes may occur in endometriosis. In the macrophage compartment, and in line with previous reports in mice^61^, we observed a significant upregulation of *IGF1* in uM2 of endometriosis cases (**Figure 5d**). In the uM1 population, a significant increase in expression of inflammatory genes *(TNFRSF1B, CEBPD)* was detected in endometriosis, in keeping with previous reports of increased inflammation in endometriosis^62,63^.

Taken together, the identified shifts in cell abundance, disease-relevant populations through fGWAS and differential gene expression analyses suggest dysregulation of stromal-immune cell homeostasis in endometriosis.

## Discussion

Globally, millions of women are affected by endometrial/uterine disorders^22–24,64^, yet the endometrium and the role of its cellular heterogeneity in these pathologies have been hugely understudied compared to other human tissues and diseases^65^. In this study, we present the HECA: the most comprehensive cellular atlas of the human endometrium assembled for individuals with/without endometriosis so far. The HECA provides a crucial step towards improving our understanding of endometrial cell heterogeneity in health and disease as it: (a) incorporates a large number of cells and individuals, (b) presents data-driven consensus cell annotation across multiple studies, (c) provides a platform for easy and rapid annotation of future scRNA-seq studies of the endometrium, and (d) enables the contextualisation of genetic association screens for endometrial/uterine disorders.

By comprehensively analysing ~614,000 high-quality cells and nuclei from 121 individuals, we substantially surpassed the number of donors and cells profiled by the initial, pioneering endometrial single-cell studies^14–21^. Having such a large sample size enabled us to identify previously unreported cell states, including a relatively rare population of *CDH2*+ (i.e. N-cadherin) epithelial cells. This population’s marker gene expression^9,34,35^, localisation within the *basalis* glands, and predicted cell-cell communication with a *basalis* fibroblast population via the *CXCR4/CXCL12* axis^36^, strongly indicated that these cells are the previously described epithelial stem/progenitor cells. Defining the transcriptomic profile of these cells opens up new avenues for exploring their role in endometrial repair and regeneration, as well as disease pathophysiology. Functional and single-cell studies that deeply characterise the seldomly sampled *basalis* layer of the endometrium (where these rare cells reside and are reported to be a heterogeneous population^2^) are now warranted in a larger number of individuals.

The HECA provides the most granular endometrial cell state annotation to date, including their spatial location in situ. Such spatial mapping was crucial for inferring the regulation and function of these cells given the spatiotemporal complexity of the endometrium. We captured multiple novel transitory cell states (e.g. preLuminal, preGlandular, subsets of decidualised stromal cells) during the early/mid secretory phase - a period crucial for endometrial receptivity preparation in response to rising progesterone levels. A tightly regulated cellular response to the changing levels of oestrogen and progesterone is essential for menstrual cycle progression, maintenance of tissue homeostasis and fertility. Thus, the newly identified cell states could present promising targets for therapy in endometrial/uterine disorders that are characterised by the disruption of hormone-dependent downstream signalling and cellular responses^66^.

Additionally, local paracrine factors and cellular crosstalk are essential for menstrual cycle progression and we provided a detailed account (and an interactive platform) of the predicted in vivo cell-cell communication across the cycle. This is an important addition to the body of existing knowledge predominantly derived from in vitro cell cultures^67–69^. Of particular interest is how TGFβ activity is controlled by various epithelial and mesenchymal cell states in both space and time. TGFβ signalling is known to suppress the activity of matrix metalloproteinases^70,71^, which are key to initiating endometrial shedding. The observed reduction in TGFβ signalling during the mid-to late secretory phase could suggest a mechanism for preparing the endometrium for shedding, or embryo implantation, and requires further studies. Interestingly, the identification and detailed description of in vivo signalling pathways involved in menstrual cycle progression could be used to refine the media used for in vitro culture of endometrial cells. For example, endometrial organoids are cultured in media supplemented with TGFβ inhibitors^72,73^, even when they are treated with hormones to mimic the menstrual cycle. Incorporating the spatial and temporal TGFβ signalling could help improve the physiological response and differentiation of these cells during the hormonal treatment, and thus eliminate some of the previously observed differences between in vivo and in vitro endometrial cells^15^.

We also revealed a range of novel interactions by which uM may aid the process of scarless endometrial regeneration, supporting previous research that proposed a role for uM in this process^74–76^. The new interactions we found pinpoint uMs pivotal roles in enhancing wound healing, boosting cellular proliferation, modulating inflammation, and stimulating angiogenesis. We also discovered that uNK1, a subset of resident NK cells which we identified to increase in proportion during the secretory phase, also expressed some angiogenic molecules, although to a lesser degree than uMs. This suggests that uMs may take on a larger role than uNKs in endometrial regeneration and angiogenesis in the non-pregnant endometrium. Interestingly, interactions between uMs and stromal cells became more evident during menstruation, emphasising the crucial role that uMs play during this phase of the cycle^77^. Understanding how the disruption of these macrophage-stromal interactions contribute to widely common menstrual disorders (e.g. abnormal uterine bleeding) could pave new paths for the development of immunology-based treatment.

Lastly, we demonstrated the utility of HECA to give cellular context to a large-scale endometriosis GWAS meta-analysis^30^. We identified two subtypes of decidualised stromal cells and macrophages as endometriosis-relevant. The observed dysregulation of stromal-immune cell homeostasis is in line with previous reports^16,20,28,29,78^, but overall, findings have been inconsistent. For example, some studies reported an increase in stromal cells in endometriosis cases, while others reported no changes. At the molecular level, our data indicated sustained WNT and dysregulated insulin signalling to be a feature of the dStromal early/mid populations in endometriosis cases. This is in line with previous observation of downregulation of *IGF2* and lack of WNT inhibition in the endometrium of women with endometriosis during the secretory phase^79–81^. At the cellular level, we previously showed that inhibition of WNT signalling by stromal cells in response to progesterone is crucial in supporting the differentiation of glandular epithelium^15^. Our current findings suggest that this process may be disrupted in endometriosis. Yet, the observed differences were subtle (i.e. exhibited small fold changes between cases and controls), requiring further validation in a larger set of samples with detailed metadata and menstrual phase annotation. To dissect the molecular pathways and validate the involvement of WNT and insulin pathways in progesterone-mediated cellular responses could now be tested using 3D in vitro models of the endometrium encompassing both stromal and epithelial cells^82^.

The HECA is a key stepping stone towards the generation of a future pan-endometrial atlas encompassing endometrial cellular heterogeneity across the lifespan and in diseases. We envision a number of endometrial/uterine atlases will be generated in the coming years, and that the HECA will guide dataset integration, cell annotation and ensure reproducibility across studies. Incorporation of datasets profiling biopsies from late secretory, peri-menstrual, menstrual and generally more finely assigned menstrual cycle phases will enrich the atlas and further improve its quality. As the atlas grows both in the number of cells and individuals profiled, it will become possible to not only look at cellular variation, but also variation at the level of individuals and link genotype to phenotype. To do so, detailed and standardised phenotypic data about the individuals studied will need to be collected (e.g. BMI, race/ethnicity, fertility status, regularity of menstrual cycles) as these factors could influence the transcriptomic profile of endometrial cells and need to be evaluated.

In summary, the HECA is the first large-scale integrated reference atlas of the human endometrium, providing a conceptual framework upon which future studies can be built. With all resources publicly available in an easy-to-access interactive format, the HECA offers a platform/tool for advancing research into endometrial physiology and disorders, as well as guiding the development of physiologically-relevant in vitro model systems of the endometrium.

## Methods

### Patient samples

Superficial endometrial samples collected for the Mareckova *et al.* dataset came from four studies: (i) Endometriosis Oxford (ENDOX), (ii) Fibroids and Endometriosis Oxford (FENOX), (iii) Sanger Human Cell Atlasing Project, and (iv) Immunology and Subfertility study. Both ENDOX (REC: 09/H0604/58) and FENOX (REC: 17/SC/0664) obtained ethical approvals from the Central University Research Ethics Committee, University of Oxford. Yorkshire & The Humber - Leeds East Research Ethics Committee approved the Sanger Human Cell Atlasing Project (REC: 19/YH/0441). The Immunology of Subfertility study (REC: 08/H0606/94) was approved by the Oxford Research Ethics Committee C. In all instances, written informed consent was provided by study participants prior to obtaining tissue samples and phenotypic data.

Full-thickness uterine wall samples were obtained from deceased transplant organ donors after ethical approval (REC: 15/EE/0152, East of England–Cambridge South Research Ethics Committee) and informed consent from the donor families. Uterus was removed within 1 h of circulatory arrest.

### Donor inclusion/exclusion criteria and endometriosis presence evaluation

Only individuals during their reproductive years were recruited and only considered having ‘natural cycles’ if they had not taken any hormonal treatment at least 3 months prior to sample collection. Donors with endometrial cancer were excluded. In addition, we aimed to exclude patients with other benign uterine/endometrial pathologies (i.e. fibroids, polyps, adenomyosis, hyperplasia). However, in some cases (*n* = 15), later histological evaluations revealed the presence of these pathologies (Supplementary Table 1). Patients taking part in the ENDOX and FENOX studies (*n* = 69) were undergoing laparoscopic surgery for suspected endometriosis or infertility reasons at the John Radcliffe Hospital, Oxford. At the beginning of surgery, a pipelle biopsy of the endometrium was taken and the presence/absence of endometriosis, including endometriosis stage (rASRM stages I-IV) assigned upon surgical evaluation during the laparoscopy. Four additional control samples (i.e. samples from donors without endometriosis) came from the Sanger Cell Atlasing Project study (*n* = 3) and Immunology of Subfertility study (*n* = 1). Absence of endometriosis was determined based on the clinical and medical history of the patients. For the Sanger Cell Atlasing Project, patients attended a coil clinic for contraceptive reasons. During the coil insertion procedure, a biopsy of the endometrium was taken in an outpatient setting. For the Immunology and Subfertility study, patients were undergoing *in vitro* fertilisation and an endometrial biopsy was taken in an outpatient setting one cycle before the patient became pregnant and had a live birth.

### Tissue processing

Superficial biopsies of the endometrium were collected using the Pipelle^®^ sampling device and immediately transferred into ice-cold phosphate buffered saline (PBS) solution (Gibco, 10010023). The endometrial tissue was then cut into smaller pieces and either moved into a cryovial and snap-frozen on dry ice (for single-nuclei extraction and processing) or moved into ice-cold HypoThermosol^®^FRS solution (Sigma-Aldrich, H4416) and stored at 4°C until further processing (either to be digested fresh or cryopreserved and digested later for single-cell processing). Where possible and sample size allowed, a small piece of tissue was also embedded in optimal cutting temperature (OCT) compound (ThermoFisher Scientific, 23730571) inside a cryomold and rapidly frozen in dry ice/isopentane slurry for histological evaluation and analyses.

Whole uterus samples used for single-cell RNA-sequencing and imaging analyses were stored in *HypoThermosoll^®^FRS at 4°C until processing*. For imaging analyses, the samples were further dissected, embedded in OCT media and rapidly frozen in dry ice/isopentane slurry. For single-cell RNA-sequencing (donor A70), to enrich endometrial cells, the endometrium was excised from the myometrium using scalpels and digested as detailed below.

### Tissue cryopreservation

Fresh tissue was cut into <1 mm^3^ segments before being resuspended with 1 ml of ice cold Cryostor solution (CS10) (C2874-Sigma). The tissue was frozen at −80°C decreasing the temperature approximately 1°C per minute. Detailed protocol available at https://www.protocols.io/view/tissue-freezing-in-cryostor-solution-processing-bgsnjwde.

### Tissue dissociation for single-cell RNA-sequencing

Cryopreserved samples were thawed at 37°C, quickly transferred to a 15 ml tube and topped-up with 13 ml of ice cold RPMI/FBS. Samples were centrifuged (500 x g, 5 min, 4°C) and the supernatant discarded. The tissue was enzymatically digested on a MACSMix rotator (set to 16 rpm speed) at 37°C in pre-warmed RPMI/FBS containing Collagenase V (Sigma-Aldrich, C9263), and DNAse I (Roche, 11284932001) with final concentrations of 1 mg/ml and 0.1 mg/ml, respectively. Digested tissue was centrifuged (500 x g, 5 min), resuspended in 10 ml of PBS and passed through a 40 µm cell strainer (BD Biosciences, 352340), generating the collagenase fraction, enriched in stromal and immune cells (Figure 3.1). The filter was back-washed with PBS into a 50 ml tube and centrifuged (500 x g, 5 min). Supernatant was discarded and any undigested tissue within the pellet was incubated with 0.25% (v/v) trypsin-EDTA (Sigma-Aldrich, T3924) and DNAse I (0.1 mg/ml) at 37°C for 15 min on a MACSMix rotator. The digestion process was stopped by adding RPMI/FBS and samples centrifuged (500 x g, 5 min). This step yielded the trypsin fraction. The collagenase fraction was centrifuged (500 x g, 5 min) and resuspended in 2 ml of red-blood-cell (RBC) lysis buffer (eBioscience, 00-4300) for 5-10 min at room temperature. After incubation, the samples were centrifuged (500 x g, 5 min), the RBC buffer discarded and both fractions (collagenase and trypsin) resuspended in 0.04% bovine serum albumin (BSA) (Sigma-Aldrich, A9418) in PBS (v/v). The generated single-cell suspensions were stored on ice and counted before being loaded separately onto the 10x Chromium chip.

In the case of two samples (donor IDs: FX1125 and FX1176), cells from the collagenase fraction were live/dead sorted prior to loading to enrich for live cells. The nuclear stain DAPI (4′,6-diamidino-2-phenylindole) was used to visualise and distinguish live/dead cells and debris.

### Tissue dissociation for single-nucleus RNA-sequencing

Snap-frozen endometrial pipelle biopsies were removed from cryovials and embedded in OCT for cryosectioning, storing them at −80°C overnight. The following day, the OCT blocks were left inside the cryostat for ~1 h to equilibrate to the chamber temperature of −20°C. The blocks were trimmed until reaching the tissue, when the first 10 µm thick sections for morphological assessment under a light microscope started to be collected. Three sections were placed on SuperFrost^®^ Plus slides (ThermoFisher, 12312148) before cutting and collecting 50 µm thick sections for nuclei extraction. Depending on tissue size, between 10 to 20 sections were placed into a 7 ml Dounce tissue grinder (Sigma-Aldrich, D9063-1SET) on dry-ice and a further three 10 µm thick sections were placed on slides and stored at −80°C for later histological staining.

Tissue collected in the Dounce tissue grinder was placed on ice inside a class II safety cabinet and incubated with 3 ml of homogenisation buffer (see Supplementary Table 5 for buffer composition) for 5 min. To help dissolve the OCT, the suspension was gently mixed with a 2 ml aspiration pipette half-way through the incubation. The tissue was then homogenised by 10-20 strokes of both pestle A and B. The number of strokes was sample-dependent - homogenisation with each pestle was performed until no resistance and tissue changes were observed. Each pestle was washed with 500 µl of the homogenisation buffer and the homogenate filtered through a 40 µm cell strainer into a new 50 ml tube. The sample was then centrifuged using the following setting: 500 x g, 6 min, 4°C, acceleration set at 0 and deceleration set to 3. After removing the supernatant, 500 µl of wash buffer (see Supplementary Table 6 for buffer composition) was added to the cell pellet and incubated for 2 min on ice. The nuclei pellet was gently resuspended using wide-bore tips to avoid damaging the nuclei, and the yield checked using a haemocytometer and trypan blue. Next, the nuclei suspension was transferred to a 1.5 ml tube and washed twice by adding 1 ml of the wash buffer and centrifugation (500 x g, 3 min, 4°C). The supernatant was removed and nuclei resuspended in 200 µl of the wash buffer (volume was nuclei yield-dependent). To remove debris and clumps, the nuclei suspension was filtered twice through the 40 µm Flowmi^®^ cell strainers and nuclei counted using a haemocytometer and trypan blue. The nuclei suspension were stored on ice until loading the 10x Chromium chip.

### Assignment of menstrual stage

OCT blocks were sectioned at 10 µM thickness and haematoxylin and eosin-stained following standard protocols. Menstrual phase was assigned based on histological evaluation by two independent pathologists. Where this was not possible, the menstrual phase was assigned based on the transcriptomic data and cellular profiles of the samples (see Supplementary Table 1).

### Donor genotyping

Buffy coats of 33 participants were genotyped using Illumina Global Screening Array (GSA) v3 with remaining genotypes retrieved from prior genotyping rounds using Affymetrix Precision Medicine Array (9 samples, including 5 in overlap with GSA v3), and Affymetrix Axiom (4 samples, 2 in overlap with Precision Medicine Array). Samples and variants quality was assessed using standard protocol^83^. Four samples were flagged (2 due to divergent ancestry, 2 due to low genotyping rate), two of which were re-genotyped on GSA v3. Variants passing QC (49.5% for Affymetrix arrays, 76.7% for GSA) were lifted from hg19 reference to GRCh38 using pyliftover and UCSC chain (v2013-12-31) with 99.92% success rate. The lifted SNPs were aligned to GRCh38.p13 reference using plink2^84^ and exported to VCF. Stand issues arising from ambiguous plink1 source data were fixed using bcftools^85^ against GRCh38.p13 reference (<5% flipped).

### Haematoxylin and Eosin (H&E) staining and imaging

Fresh frozen sections were removed from −80°C storage and air dried before being fixed in 10% neutral buffered formalin for 5 minutes. After rinsing with deionised water, slides were dipped in Mayer’s Haematoxylin solution for 90 seconds. Slides were completely rinsed in 4-5 washes of deionised water, which also served to blue the haematoxylin. Aqueous eosin (1%) was manually applied onto sections with a pipette and rinsed with deionised water after 1-3 seconds. Slides were dehydrated through an ethanol series (70%, 70%, 100%, 100%) and cleared twice in 100% xylene. Slides were coverslipped and allowed to air dry before being imaged on a Hamamatsu Nanozoomer 2.0HT digital slide scanner.

### Multiplexed smFISH and high-resolution imaging

Large tissue section staining and fluorescent imaging was conducted largely as described previously^86^. Sections were cut from fresh frozen embedded in OCT at a thickness of 10 μm using a cryostat, placed onto SuperFrost Plus slides (VWR) and stored at −80°C until stained. Tissue sections were then processed using a Leica BOND RX to automate staining with the RNAscope Multiplex Fluorescent Reagent Kit v2 Assay (Advanced Cell Diagnostics, Bio-Techne), according to the manufacturers’ instructions. Probes used are found in Supplementary Table 7. Prior to staining, tissue sections were post-fixed in 4% paraformaldehyde in PBS for 15 minutes at 4°C, then dehydrated through a series of 50%, 70%, 100%, and 100% ethanol, for 5 minutes each. Following manual pre-treatment, automated processing included epitope retrieval by protease digestion with Protease IV for 30 minutes prior to probe hybridisation. Tyramide signal amplification with Opal 520, Opal 570, and Opal 650 (Akoya Biosciences) and TSA-biotin (TSA Plus Biotin Kit, Perkin Elmer) and streptavidin-conjugated Atto 425 (Sigma Aldrich) was used to develop RNAscope probe channels. Stained sections were imaged with a Perkin Elmer Opera Phenix High-Content Screening System, in confocal mode with 1 μm z-step size, using a 20X (NA 0.16, 0.299 μm/pixel); 40X (NA 1.1, 0.149 μm/pixel); water-immersion objective. Channels: DAPI (excitation 375 nm, emission 435-480 nm), Atto 425 (ex. 425 nm, em. 463-501 nm), Opal 520 (ex. 488 nm, em. 500-550 nm), Opal 570 (ex. 561 nm, em. 570-630 nm), Opal 650 (ex. 640 nm, em. 650-760 nm). *Image stitching:* Confocal image stacks were stitched as two-dimensional maximum intensity projections using proprietary Acapella scripts provided by Perkin Elmer.

### 10x Genomics Chromium GEX library preparation and sequencing

Both cells and nuclei undergoing scRNA-seq and snRNA-seq were loaded according to the manufacturer’s protocol for the Chromium Single Cell 3′ Kit v.3.0, and v3.1 (10X Genomics) to attain between 2,000 and 10,000 cells/nuclei per reaction. Libraries were sequenced, aiming at a minimum coverage of 50,000 raw reads per cell, on the Illumina Novaseq 6000 system; using the sequencing format; read 1: 28 cycles; i7 index: 10 cycles, i5 index: 10 cycles; read 2: 90 cycles.

### 10x Genomics Visium library preparation and sequencing

We generated 10x Genomics Visium transcriptomic slides from two superficial biopsies. Briefly, 10 micron cryosections were cut and placed on Visium slides v1 3’. These were processed according to the manufacturer’s instructions. Briefly, sections were fixed with cold methanol, stained with haematoxylin and eosin and imaged on a Hamamatsu NanoZoomer S60 before permeabilisation, reverse transcription and cDNA synthesis using a template-switching protocol. Second-strand cDNA was liberated from the slide and single-indexed libraries prepared using a 10x Genomics PCR-based protocol. Libraries were pooled and sequenced on a Novaseq 6000), with the following sequencing format; read 1: 28 cycles, i7 index: 10 cycles, i5 index: 10 cycles and read 2: 90 cycles.

### External human endometrial scRNA-seq and Visium datasets

We collected raw sequencing data from previously published human endometrial scRNA-seq datasets. Specifically, we downloaded publicly available .fastq files either from Gene Expression Omnibus (GEO) or ArrayExpress. These datasets included: (i) Wang et al. (GEO accession number GSE111976), (ii) Garcia-Alonso et al. (ArrayExpress accession number E-MTAB-10287), (iii) Tan et al. (GEO accession number GSE179640), (iv) Lai et al. (GEO accession number GSE183837), (v) Fonseca et al. GEO accession number GSE213216), and (vi) Huang et al. (GEO accession number GSE214411).

For spatial transcriptomics analysis, we used the 10x Genomics Visium from two full thickness uterus previously generated by us, available at ArrayExpress (accession number E-MTAB-9260).

### Alignment and quantification of sc/snRNA-seq data

Reads from both the newly generated scRNA-seq/snRNA-seq libraries and external datasets were alignment to the 10x Genomics’ human reference genome GRCh38-2020-A, followed by cell calling, transcript quantification and quality control (QC) using the Cell Ranger Software (version 6.0.2; 10X Genomics) with default parameters. Cell Ranger filtered count matrices were used for downstream analysis.

### Downstream sc/snRNA-seq analysis

#### Donor demultiplexing and doublet identification

For 84 of the newly generated libraries (26 in the scRNA-seq and 58 in the snRNA-seq datasets) we multiplexed cell suspensions from two different donors. To ensure that we could confidently assign cells back to their donor, we genotyped some donors as described in the *Donor genotyping* section above, and then pooled sample combinations in a way that each scRNA-seq/snRNA-seq library contained at least one genotyped donor.

To assign each cell/nuclei in the scRNA-seq/snRNA-seq libraries back to their donor-of-origin, we genotyped each barcode. Specifically, we called the SNPs in the reads from each barcode and piled them up using the cellSNP tool v1.2.2. Here, reads were genotyped from the Cell Ranger BAM files using a reference list of human common variants from the 1000 Genome Project (hg38 version with minor allele frequency (MAF) > 0.0005) that we downloaded from https://sourceforge.net/projects/cellsnp/files/SNPlist. Once the cells in scRNA-seq/snRNA-seq libraries were genotyped, we linked them back to their donor-of-origin genotype (obtained using Illumina Global Array) using vireoSNP v0.5.8 with default parameters (n_donor = 2). Barcodes classified as either “doublet” (i.e. containing the two genotypes) or “unassigned” were discarded in downstream analysis.

#### Doublet detection based on transcriptional mixtures

We quantified cell-doublet likelihood for each barcode with Scrublet software on a per-library basis. We used a two-step diffusion doublet identification followed by Bonferroni-FDR correction and a significance threshold of 0.01, as described in ^87^. Barcodes estimated as doublets were not excluded from the initial analysis, instead these were kept in the downstream analysis and used to identify doublet-enriched clusters.

#### Quality filters, batch correction and clustering

For both scRNA-seq and snRNA-seq libraries, we used the filtered count matrices from Cell Ranger 6.0.2 for downstream analysis and analysed them with Scanpy v.1.7.0, with the pipeline following their recommended standard practises. We applied stringent QC to further filter the cells called by Cell Ranger to retain only high-quality cells. Specifically, we excluded cells either (i) expressing fewer than 1,000 genes or (ii) with a mitochondrial content higher than 20%. For some datasets, these filters discarded more than 50% of the initial called cells.

Next, we flagged cell-cycle genes using a data-driven approach as described in ^87,88^. To do so, after converting the expression space to log(CPM/100 + 1), we transpose the object to gene space, performing PCA, neighbour identification and Leiden clustering. The gene members of the gene cluster encompassing well-known cycling genes (*CDK1*, *MKI67*, *CCNB2* and *PCNA*) were all flagged as cell cycling genes, and discarded in each downstream analysis. In parallel, we also used the scanpy function “score_genes_cell_cycle” to infer the cell cycle stage of each cell (i.e. G1, G2/M or S) that was later used to interpret the clusters.

Next, we generated an integrated manifold for scRNA-seq and snRNA-seq datasets separately. The scRNA-seq manifold included data from 6 previously published studies as well as the scRNA-seq data newly generated by us. The snRNA-seq exclusively contains newly generated data for this study. To minimise cell cycle bias, the previously flagged cell-cycle genes were excluded. The integrated manifolds were generated using single-cell Variational Inference (scVI) v0.6.8, with both the donor and study id (for scRNA-seq only) as batches. All the remaining parameters were kept as default, with n_latent= 32, n_layers=2. The scVI low dimensional space was estimated on the top 2,000 most highly variable genes in each dataset, which were defined using *Seurat v3* flavour on the raw counts. With the resulting scVI-corrected latent representation of each cell, we estimated the neighbour graph, generated a Uniform Manifold Approximation and Projection (UMAP) visualisation and performed Leiden clustering.

The same strategy was used to zoom-in into each of the four main cell lineages (i.e. epithelial, mesenchymal, immune and endothelial) to further resolve the cellular heterogeneity in those compartments. Here, we subset the cells to those in the lineage and repeated scVI integration using the top 2,000 most highly variable genes within each lineage. The donor and the study id were kept as batches, with default parameters, n_latent= 64 and n_layers=2. For the zoom-in analysis into the immune compartment, donors taking exogenous hormones (Tan et al dataset) were excluded due to integration challenges.

### Annotation of cell types

We performed a full re-annotation of the cell clusters in the integrated scRNA-seq manifold. First, we carried out a new quality control round to exclude clusters that are likely driven by technical artefacts (i.e. low QC cells or doublets). Briefly, we flagged as *low QC clusters* those that (i) express an overall lower number of genes, (ii) express an overall lower number of counts, (iii) display a higher than average mitochondrial or nuclear RNA content and, importantly (iv) do not express any distinctive gene (and thus are not representing any independent biological entity). Next, we flagged as *doublets* those clusters that met the following criteria (i) exhibit higher scrublet doublet score; (ii) express marker genes from multiple lineages (for example, display both epithelial and immune markers) and (iii) do not express any distinctive gene. Distinctive marker genes were identified using TF-IDF, as implemented in the SoupX package v.1.5.0.

To assign cell type labels to remaining high-quality clusters, we took into account the following variables: (i) the menstrual cycle phase bias (or any other clinical variable such exogenous hormones, endometriosis, etc), (ii) the expression of previously described markers, (iii) the differentially expressed genes and (iv) the spatial location, either by performing smFISH or by deconvoluting the cellular composition of Visium spots.

Because of the higher gene coverage of the scRNA-seq data, cell type identification and annotation was done primarily on the integrated scRNA-seq dataset. To annotate the snRNA-seq clusters, we trained a Support Vector Machine (SVM) classifier (sklearn.svm.SVC) on the scRNA-seq dataset and transferred labels onto the denoised (i.e. decontaminated of ambient RNA) snRNA-seq dataset. Denoising of snRNA-seq was done with DecontX from the R celda package v1.6.1. Predicted cell type annotations on snRNA-seq were validated or disproved by looking at the expression of marker genes.

### Query-to-HECA mapping

We used the scArches model surgery framework^32^ to project new samples onto the same latent space as single-cell HECA. The scVI model used in main analysis was trained using both dataset and sample ID as covariates. In order to build a model compatible with the scArches framework, we trained a scANVI model with only sample ID as batch covariate. We trained the reference scANVI model for 20 epochs, based on an scVI model with n_layers = 2. The surgery model was trained for 100 epochs with weight_decay=0.0 to ensure reference cell embeddings would remain identical. To obtain joint embeddings, we concatenated gene expression counts from HECA reference cells and query samples into a single object and used the surgery encoder to get latent representations. We then computed the kNN graph (default parameters) and UMAP (min_dist = 0.4) on the joint embeddings. We evaluated the quality of the query to reference mapping by examining the alignment on the UMAP and the concordance of marker gene expression in HECA reference cells and query samples.

We provide step-by-step scArches tutorials at https://github.com/ventolab/HECA-Human-Endometrial-Cell-Atlas/blob/main/tutorials/query_to_ref_mapping.ipynb to support mapping new samples to HECA reference cells based on any input gene expression count matrix.

### Alignment and quantification of Visium data

The newly generated 10x Visium spatial sequencing data was processed using Space Ranger Software (v.2.0.1) to identify the spots under tissue, align reads to the 10x Genomics’ human reference genome GRCh38-2020-A and quantify gene counts. Spots were automatically aligned to the paired H&E images by Space Ranger software. All spots under tissue detected by Space Ranger were included in downstream analysis.

### Downstream analysis of Visium data

#### Location of cell types in Visium data

We spatially map the cell types from the scRNA-seq dataset on the Visium slides with cell2location tool v0.06-alpha^89^. We deconvoluted both, the Visium slides newly generated in this study from superficial biopsies and the ones downloaded from E-MTAB-9260 covering full thickness uterus. As reference, we used the cell type signatures from the scRNA-seq dataset, subsetting the cells to those expressing more than 2,000 genes. Cell2location was run with default parameters, with the exception of *cells_per_spot* which was set to 20. Each Visium section was analysed separately. The estimated abundance for each cell type was visualised following the *cell2location* tutorial.

### Cell-cell communication analysis with CellPhoneDB

Because two cell types can only interact paracrinally or juxtacrinally if they co-localise in space and time, we first manually classified the cell types into the spatiotemporal microenvironments where these coexists (for example, endothelial and PV cells coexist in the vessels, while preGlandular coexists with dStromal early cells in the *functionalis* layer of the early secretory endometrium). Spatial location was derived from prior knowledge, smFISH experiments or cell type deconvolution of Visium spots with cell2location. The temporal location was directly derived from the menstrual phase where the cell types are detected.

To identify paracrine or juxtacrine interactions between the cells co-localising in an endometrial microenvironment, we used the DEGs-based method of CellphoneDB v4.1^90^. Using this method, we retrieved interacting pairs of ligands and receptors meeting the following requirements: i) all the interacting partners were expressed by at least 10% of the cell type under consideration; ii) the interacting cell type pairs share an endometrial microenvironment and iii) at least one of the interacting partners (for example, either the ligand or the receptor) was significantly upregulated in the corresponding cell type (Wilcoxon Tests; adjusted p-value < 0.01 and a log2 fold change > 0.75). Differential expression analysis was performed on a per-lineage approach to identify the genes specifically upregulated in a cell state, compared to the other cell states in the same lineage. Donors under exogenous hormonal therapy were excluded from the analysis.

### Differential cell abundance (DCA)

To quantify changes in cellular composition, we used differential abundance analysis on cell neighbourhoods with RMilo v1.6.0^91^.

To evaluate compositional changes of immune cells between the proliferative or the secretory phases of the menstrual cycle, we first calculated the KNN graph derived from the scVI immune-embedding subsetted to contain only superficial biopsies from controls (excluding donors under exogenous hormone therapy). Next, we assigned cells to neighbourhoods and counted the number of cells belonging to each cell type in each neighbourhood. We assigned each neighbourhood to a cell type label based on majority voting of the cells belonging to that neighbourhood. Cell neighbourhoods where less than 70% of cells came from a single cell type were labelled as “Mixed neighbourhoods” and discarded. To test for differential abundance across the menstrual cycle, we divided the samples into proliferative and the secretory phases. RMilo models the cell count in neighbourhoods as a negative binomial generalised linear model, using a log-linear model to model the effects of menstrual phase on cell counts, while accounting for the total number of cells over all the neighbourhoods. When analysing the scRNA-seq dataset, we also included the study id as covariate of the model to account for the variability between laboratory/technical batches. A neighbourhood was associated with the proliferative phase if SpatialFDR < 0.1 and logFC < 0, or the secretory phase if SpatialFDR < 0.1, logFC > 0. The analysis was performed primarily with the snRNA-seq datasets to minimise laboratory bias, and validated on the scRNA-seq dataset where we included the study id as model covariate.

To evaluate compositional changes of mesenchymal, epithelial, endothelial and immune cells between endometriosis and controls, we again relied on the KNN graph derived from the scVI lineage-subanalysis embedding. Nuclei were subsetted to those coming only from superficial biopsies, excluding donors under exogenous hormone therapy. Following the strategy described above, we estimated differential abundance between control and endometriosis case samples using the snRNA-seq dataset, as it has an even coverage of cases and controls along the menstrual cycle and avoids laboratory bias. Stromal and epithelial populations are menstrual-phase specific, and were tested considering donors in the corresponding phase. For testing differences in immune cells we instead added the menstrual phase as a covariate in the model. A neighbourhood was associated with control samples if SpatialFDR < 0.1 and logFC < 0, or endometriosis case samples if SpatialFDR < 0.1, logFC > 0.

### Cell type enrichment analysis for Endometriosis-GWAS genes

To study the association between the endometrial cell populations in our atlas and previously identified endometriosis GWAS loci, we used the functional GWAS (fGWAS) approach described in^92^. This approach evaluates the enrichment of various functional annotations for molecular quantitative traits (in this case, the gene expression signature of a cell type) and GWAS loci (in this case, the cis-regulatory variants associated with endometriosis). Here, genetic variants were linked to genes if they map to their cis-regulatory region, which is defined as ±500 Kb centred at the transcription start site (TSS) of the gene. The association statistics (the log odds ratios and standard errors) were transformed into the approximate Bayes factors using the Wakefield method^92,93^. The Bayes factors of variants mapping to each gene cis-regulatory region were weighted and averaged by the prior probability, estimated as the exponential function to TSS proximity. Finally, the enrichment of each cell type was estimated as the maximum likelihood estimator of the effect size for the cell-type-specific expression.

Endometriosis GWAS loci were derived from the full summary statistics of our recent endometriosis GWAS meta-analysis^30,94^ excluding the *23andMe* dataset. The full summary statistics, indicating the SNP position, beta value and standard error used to perform the fGWAS analysis, are publicly available from EBI GWAS Catalog (GCST90205183).

### Differential gene expression endometriosis vs controls

We evaluated the magnitude and significance of the differences in gene expression between endometriosis patients and controls using limma v.3.54.2. First, to avoid unwanted confounding effects, we subsetted the data to contain only superficial biopsies and excluded donors under exogenous hormonal therapy. Secondly, to account for within-sample correlations (i.e. cells coming from the same donor), pseudobulking with sum aggregation was performed prior to applying limma. Briefly, we generated 3 pseudobulks per donor and per cell type by aggregating the cells of each cell type and taking the mean gene expression within the cell type. Finally, we tested for differential expression between conditions (endometriosis vs control) using the limma-voom approach. The analysis was performed on the scRNA-seq datasets, and we reported as differentially expressed genes with FDR < 0.1.

## Data availability

Datasets are available from ArrayExpress (www.ebi.ac.uk/arrayexpress), with accession number pending. Multiplexed smFISH images are available from BioStudies (www.ebi.ac.uk/biostudies), with accession number pending. All data is public access. Source data are provided with this paper. scRNA-seq and snRNA-seq datasets to reproduce UMAPs and dotplots can be accessed and downloaded through the web portals https://www.reproductivecellatlas.org/endometrium_reference.html.

## Code availability

All the code used for data analysis is available at: https://github.com/ventolab/HECA-Human-Endometrial-Cell-Atlas.

## Acknowledgements

This publication is part of the Human Cell Atlas *–* www.humancellatlas.org/publications/. The authors would like to thank the participants of the ENDOX and FENOX studies in the Oxford Endometriosis CaRe Centre for donating samples and data. We are also grateful to the transplant organ donors and their families for the samples donated through the Cambridge Biorepository for Translational Medicine. We would also like to thank Kelly Barrett, Carol Hubbard and Lisa Buck for patient recruitment and clinical sample collection; the Sanger Cellular Generation and Phenotyping (CGaP) Core Facility, Sanger Core Sequencing pipeline for support with sample processing and sequencing library preparation; Martin Prete and Simon Murray for insightful comments and web portal support; Tarryn Porter and the Cellular Genetics wet lab team for experimental support; Antonio García from Bio-Graphics for scientific illustrations; Aidan Maartens for proofreading.

## Funding

This research was funded in part, by the Wellcome Trust Grant 206194 and 220540/Z/20/A and 203141/Z/16/Z; the European Union’s Horizon 2020 research and innovation programme HUTER under grant agreement No 874867; grant number 2022-249429(5022) from the Chan Zuckerberg Foundation; and the John Fell Fund from the University of Oxford. M.Ma is funded by the Medical Research Council. Sample collection at Imperial was supported by Borne, grant no P84654.

## Author information

R.V.T, M.Ma, and L.G-A conceived and designed the experiments and analyses. L.G-A analysed the data with contributions from M.Ma, M.Mo, V.L, S.H, M.K and M.L. M.Ma, C.S.S. and A.O. performed sample processing. C.I.M performed the imaging experiments. L.G-A and R.P developed the cell-cell communication platform. K.Ga oversaw patient metadata collection for ENDOX and FENOX studies. Ke.G and S.Y performed menstrual cycle staging. E.V.W, V.M, K.T.M, K.S-P and I.G. collected some of the samples analysed. M.Ma, L.G-A and R.V.T. interpreted the data with contributions from M.Mo, V.L, and S.H. M.Ma, L.G-A and R.V.T wrote the manuscript. R.V-T and K.T.Z supervised the work with contributions from C.M.B, R.A.D and J.S. All authors read and approved the manuscript.

## Competing interests

K.T.Z and C.M.B have received grant funding from Bayer AG, AbbVie Inc., Roche Diagnostics Inc., Volition Rx, MDNA Life Sciences, and Precision Life, unrelated to the work presented in this paper. K.T.Z is also a Board member of the World Endometriosis Research Foundation. The remaining authors declare no competing interests.

## Supplementary Material

### Supplementary Figures

**Supplementary Figure 1. Single-cell RNA-sequencing datasets of the Human Endometrial Cell Atlas (HECA) and the cervix. a,** UMAP projections of scRNA-seq data for HECA colured by cell lineage, dataset, menstrual cycle group, cell cycle phase and biopsy type. **b,** Dot plot showing normalised, log-transformed and variance-scaled expression of genes (*x*-axis) characteristic of the main cell lineage (*y*-axis) in the HECA. **c,** Dot plot showing normalised, log-transformed and variance-scaled expression of genes (*x*-axis) characteristic of a selection of mesenchymal and endothelial cells (*y*-axis) in the HECA. **d,** Bar plot showing the cellular composition of endometrial biopsies belonging to the different menstrual cycle groups (*y-*axis). **e,** UMAP projection of a scANVI representation of the HECA coloured by the cell states identified. The MUC5B, KRT5 and HOXA13 populations are outlined by red dotted-lined shapes. **f,** UMAP projection of the Liu et al. 2023 scRNA-seq dataset of the human cervix coloured by louvain clusters and their correspondence to the four main cell lineages (epithelial, endothelial, mesenchymal and immune). **g,** Dot plot showing normalised, log-transformed and variance-scaled expression of genes (*x*-axis) characteristic of the cell clusters identified in the Liu et al. 2023 cervix dataset (*y*-axis). Highlighted by purple rectangles are the epithelial and mesenchymal clusters that expressed markers characteristic of the MUC5B, KRT5 and HOXA13 cell populations defined in the HECA. **h,** UMAP projection of the mapping of the Liu et al. 2023 cervix dataset onto the scANVI representation of the HECA coloured by the cell states identified in the HECA and the Liu et al. 2023 cervix dataset (dark grey). The MUC5B, KRT5 and HOXA13 populations of the HECA are outlined by red dotted-lined shapes. **i,** UMAP projection of the mapping of the Liu et al. 2023 cervix dataset onto the scANVI representation of the HECA coloured by the cell clusters identified in the Liu et al. 2023 cervix dataset. The MUC5B, KRT5 and HOXA13 populations of the HECA are outlined by red dotted-lined shapes. dStromal, decidualised stromal cells; ePV, endometrial perivascular cells; eStromal, endometrial stromal cells specific to proliferative phase; HECA, human endometrial cell atlas; MMPs, matrix metalloproteinases; NK, natural killer cells; scRNA-seq, single-cell RNA-sequencing; scANVI, single-cell ANnotation using Variational Inference; T, T cells; UMAP, uniform manifold approximation and projection; uSMCs, uterine smooth muscle cells.

**Supplementary Figure 2. Single-nucleus RNA-sequencing cell state identification and marker gene expression. a,** UMAP projections of the snRNA-seq data coloured by cell lineage, cell cycle phase, menstrual cycle group, and endometriosis status. **b,** UMAP projections of the epithelial cell lineage of the snRNA-seq dataset coloured by the identified epithelial cell states of the HECA as assigned by label transfer. **c,** UMAP projections of the mesenchymal cell lineage of the snRNA-seq dataset coloured by the identified mesenchymal cell states of the HECA as assigned by label transfer. **d,** Dot plot showing normalised, log-transformed and variance-scaled expression of genes (*x*-axis) characteristic of the endothelial and immune nuclei (*y*-axis). **e,** Bar plot showing the cellular composition of endometrial biopsies belonging to the different menstrual cycle groups (*y-*axis). **f,** Dot plot showing normalised, log-transformed and variance-scaled expression of genes (*x*-axis) characteristic of the identified epithelial cell states (*y*-axis) in snRNA-seq data. **g,** Dot plot showing normalised, log-transformed and variance-scaled expression of genes (*x*-axis) characteristic of the identified mesenchymal cell states (*y*-axis) in snRNA-seq data. dStromal, decidualised stromal cells; ePV, endometrial perivascular cells; eStromal, endometrial stromal cells specific to proliferative phase; HECA, human endometrial cell atlas; MMPs, matrix metalloproteinases; mPV, myometrial perivascular cells; Prolif., proliferative; secret., secretory; snRNA-seq, single-nucleus RNA-sequencing; UMAP, uniform manifold approximation and projection; uSMCs, uterine smooth muscle cells.

**Supplementary Figure 3. Cellular heterogeneity of samples from donors taking exogenous hormones in scRNA-seq and snRNA-seq data. a,** UMAP projections of the scRNA-seq data coloured by hormonal treatment taken. **b,** Overview of the number of donors and cells per hormonal treatment taken in each dataset profiled by scRNA-seq. **c,** Bar plot showing the cellular composition of endometrial biopsies from donors taking the different hormonal treatment (*y*-axis) in the scRNA-seq data. **d,** UMAP projections of the snRNA-seq data coloured by hormonal treatment taken. **e,** Overview of the number of donors and cells per hormonal treatment taken profiled by snRNA-seq. **c,** Bar plot showing the cellular composition of endometrial biopsies from donors taking the different hormonal treatment (*y*-axis) in the snRNA-seq dataset. dStromal, decidualised stromal cells; ePV, endometrial perivascular cells; eStromal, endometrial stromal cells specific to proliferative phase; MMPs, matrix metalloproteinases; mPV, myometrial perivascular cells; Prolif., proliferative; scRNA-seq; single-cell RNA-sequencing; secret., secretory; snRNA-seq, single-nucleus RNA-sequencing; UMAP, uniform manifold approximation and projection; uSMCs, uterine smooth muscle cells.

**Supplementary Figure 4. Spatial transcriptomics mapping of epithelial cell populations. a,** Visium spatial transcriptomics data and an H&E image of the same tissue section are shown. Spot colour indicates estimated cell state density for a specific population in each Visium spot, as computed by cell2location. Spatial mapping of the SOX9 functionalis I (CDH2+) and SOX9 functionalis II populations is shown in a section of a whole-uterus biopsy from donor A13. **b,** Visium spatial transcriptomics data and an H&E image of the same tissue section are shown. Spot colour indicates estimated cell state density for a specific population in each Visium spot, as computed by cell2location. Spatial mapping of the preLuminal, Luminal, preGlandular and Glandular populations is visualised in a section of a superficial biopsy from donor FX0033 (early secretory phase). **c,** High-resolution multiplexed smFISH of a section of a superficial biopsy from donor FX9006 (early secretory phase) showing the expression of *DAPI* (white, nuclei), *EPCAM* (magenta, epithelial cells), *CBR3* (cyan, preGlandular cells), and *OPRK1* (yellow, preGlandular cells). The dashed outline indicates the area shown magnified to the right. The magnified image shows the glandular region with high and co-localised *OPRK1* and *CBR3* signal. White arrows indicate luminal regions with low *OPRK1* and *CBR3* signal. **d,** High-resolution multiplexed smFISH of full thickness endometrium sections from the proliferative phase (donors A66 and A13) and secretory phase (donor A30) showing the expression of *DAPI* (white, nuclei), *EPCAM* (magenta, epithelial cells), and *MUC5B* (yellow, epithelial cells). For each panel, the dashed outline indicates the area shown magnified. Asterisks indicate some of the regions where the MUC5B signal was detected and varied across samples. Scale bars are 100 µm, unless differently specified. smFISH, single molecule fluorescence in situ hybridisation.

**Supplementary Figure 5. Expression of receptors involved in TGFβ, insulin, retinoic acid and WNT signalling.** Dot plot showing normalised, log-transformed and variance-scaled expression of genes coding for TGFβ, insulin, retinoic acid and WNT signalling receptors (*x*-axis) in the epithelial and mesenchymal cell states identified (*y*-axis) in the scRNA-seq data. eStromal, endometrial stromal cells specific to proliferative phase; dStromal, decidualised stromal cells; MMPs, matrix metalloproteinases; scRNA-seq, single-cell RNA-sequencing; TGFβ, transforming growth factor beta; uSMCs, uterine smooth muscle cells.

**Supplementary Figure 6. Immune cells in scRNA-seq and snRNA-seq data. a,** UMAP projections of scRNA-seq data for immune cells coloured by dataset, menstrual cycle group, cell cycle phase and biopsy type. **b,** UMAP projections of snRNA-seq data for immune cells coloured by menstrual cycle group and cell cycle phase. **c,** UMAP projection of snRNA-seq data for immune cells coloured by the probability of assigning the immune cell types identified in the scRNA-seq data. Support Vector Machine (SVM) classifier was trained using the immune cell scRNA-seq data and the predicted cell type annotations were then projected onto the snRNA-seq data with the probability shown. **d,** Dot plot showing normalised, log-transformed and variance-scaled expression of genes (*x*-axis) characteristic of the identified immune cell states (*y*-axis) in the scRNA-seq data. **e,** Dot plot showing normalised, log-transformed and variance-scaled expression of genes (*x*-axis) characteristic of the identified immune cell states (*y*-axis) in the snRNA-seq data. **f,** Beeswarm plot of the distribution of log fold change across the menstrual cycle (proliferative and secretory phases) in neighbourhoods containing immune cells from different cell type clusters in snRNA-seq data. Differentially abundant neighbourhoods at log fold change > 2.5 and spatial FDR < 0.1 are coloured. **g,** Visium spatial transcriptomics data for donors A13 (proliferative phase) and A30 (secretory phase) are shown. Spot colour indicates estimated cell state density for a specific population of perivascular cells (mPV, ePV-1a, ePV-1b and ePV-2) in each Visium spot, as computed by cell2location. **h,** Dot plot showing normalised, log-transformed and variance-scaled expression of genes (*x*-axis) characteristic of the identified endothelial, perivascular and stromal cells (*y*-axis) in the scRNA-seq data. cDC, conventional dendritic cells; eStromal, endometrial stromal cells specific to proliferative phase; ePV, endometrial perivascular cells; FDR, false discovery rate; ILC3, innate lymphoid cell type 3; mPV, myometrial perivascular cells; pDC, plasmacytoid dendritic cells; RBC, red blood cells; scRNA-seq, single-cell RNA-sequencing; snRNA-seq, single-nucleus RNA-sequencing; SVM, support vector machine; T Reg, T regulatory cells; uM, uterine macrophages; UMAP, uniform manifold approximation and projection; uNK, uterine natural killer cells.

**Supplementary Figure 7. Predicted cell-cell interactions underpinning endometrial regeneration and angiogenesis. a,** Dotplot plot reporting the variance-scaled mean expression of the two or more (if heteromeric complexes) transcripts coding for the interacting proteins in pairs of cell types. Red circles indicate that at least one of the interacting partners is differentially expressed in one of the cell types in the pair. Interactions are classified based on whether they are predicted to play a role in recruitment, wound healing or immunomodulation during endometrial regeneration. **b,** Dotplot plot reporting the variance-scaled mean expression of the two or more (if heteromeric complexes) transcripts coding for the interacting proteins in pairs of cell types. Red circles indicate that at least one of the interacting partners is differentially expressed in one of the cell types in the pair. Interactions are classified based on whether they are predicted to play a role in cell recruitment or pro-angiogenic processes within the vascular niche.

### Supplementary Tables

**Supplementary Table 1:** Harmonised metadata of samples analysed.

**Supplementary Table 2:** CellRanger QC outputs for all newly generated data.

**Supplementary Table 3:** Differentially expressed genes reported for stromal cells.

**Supplementary Table 4:** Differentially expressed genes reported for macrophages.

**Supplementary Table 5:** Reagents used for the snRNA-seq homogenisation buffer.

**Supplementary Table 6:** Reagents used for the snRNA-seq wash buffer.

**Supplementary Table 7:** List of smFISH probes used for smFISH imagining.

