## Supplementary Figures for "An integrated single-cell reference atlas of the human endometrium"

### Supplementary Figure 1

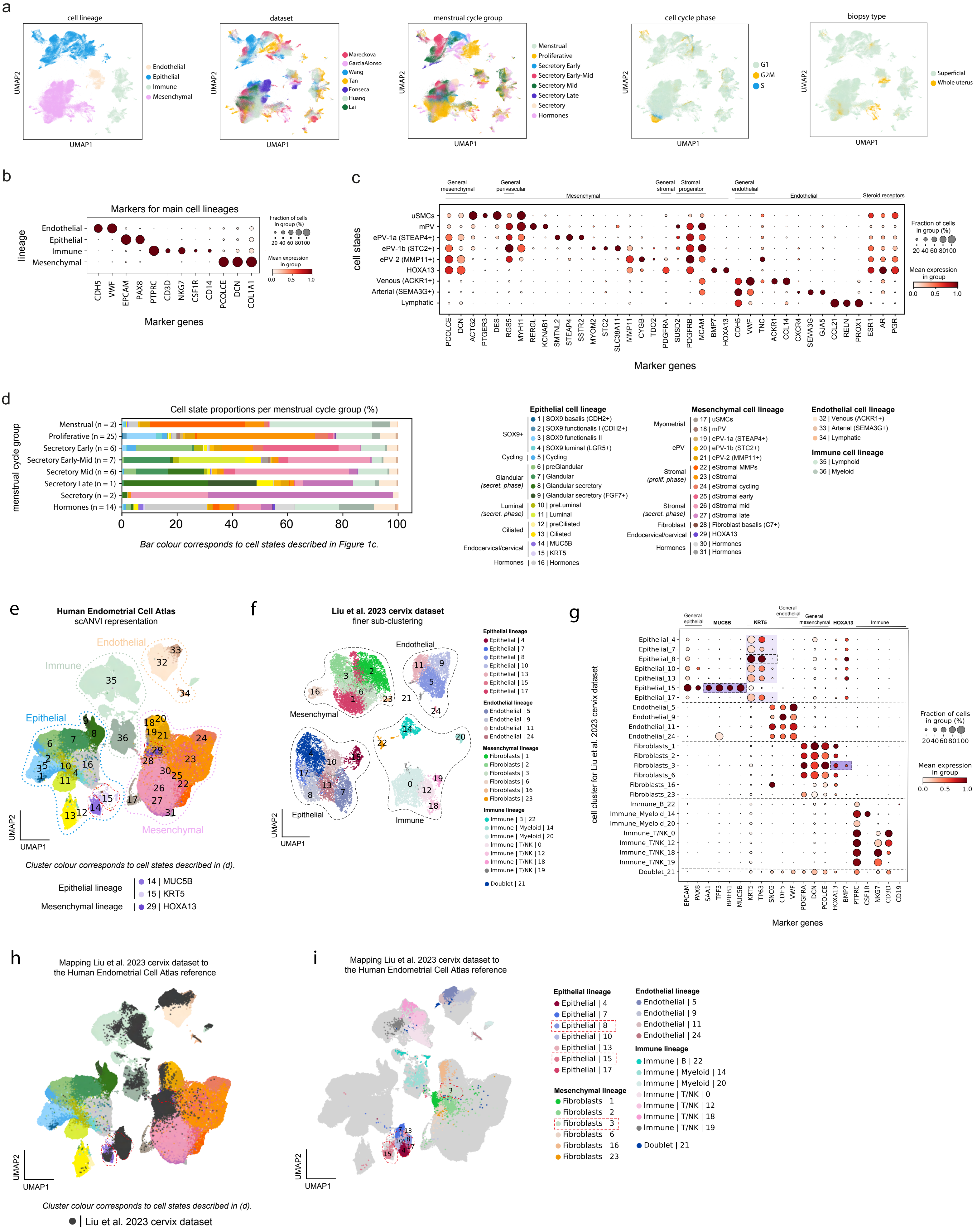

**Supplementary Figure 1. Single-cell RNA-sequencing datasets of the Human Endometrial Cell Atlas (HECA) and the cervix. a,** UMAP projections of scRNA-seq data for HECA coloured by cell lineage, dataset, menstrual cycle group, cell cycle phase and biopsy type. **b,** Dot plot showing normalised, log-transformed and variance-scaled expression of genes (*x*-axis) characteristic of the main cell lineage (*y*-axis) in the HECA. **c,** Dot plot showing normalised, log-transformed and variance-scaled expression of genes (*x*-axis) characteristic of a selection of mesenchymal and endothelial cells (*y*-axis) in the HECA. **d,** Bar plot showing the cellular composition of endometrial biopsies belonging to the different menstrual cycle groups (*y*-axis). **e,** UMAP projection of a scANVI representation of the HECA coloured by the cell states identified. The MUC5B, KRT5 and HOXA13 populations are outlined by red dotted-lined shapes. **f,** UMAP projection of the Liu et al. 2023 scRNA-seq dataset of the human cervix coloured by louvain clusters and their correspondence to the four main cell lineages (epithelial, endothelial, mesenchymal and immune). **g,** Dot plot showing normalised, log-transformed and variance-scaled expression of genes (*x*-axis) characteristic of the cell clusters identified in the Liu et al. 2023 cervix dataset (*y*-axis). Highlighted by purple rectangles are the epithelial and mesenchymal clusters that expressed markers characteristic of the MUC5B, KRT5 and HOXA13 cell populations defined in the HECA. **h,** UMAP projection of the mapping of the Liu et al. 2023 cervix dataset onto the scANVI representation of the HECA coloured by the cell states identified in the HECA and the Liu et al. 2023 cervix dataset (dark grey). The MUC5B, KRT5 and HOXA13 populations of the HECA are outlined by red dotted-lined shapes. **i,** UMAP projection of the mapping of the Liu et al. 2023 cervix dataset onto the scANVI representation of the HECA coloured by the cell clusters identified in the Liu et al. 2023 cervix dataset. The MUC5B, KRT5 and HOXA13 populations of the HECA are outlined by red dotted-lined shapes. dStromal, decidualised stromal cells; ePV, endometrial perivascular cells; eStromal, endometrial stromal cells specific to proliferative phase; HECA, human endometrial cell atlas; MMPs, matrix metalloproteinases; NK, natural killer cells; scRNA-seq, single-cell RNA-sequencing; scANVI, single-cell ANnotation using Variational Inference; T, T cells; UMAP, uniform manifold approximation and projection; uSMCs, uterine smooth muscle cells.

### Supplementary Figure 2

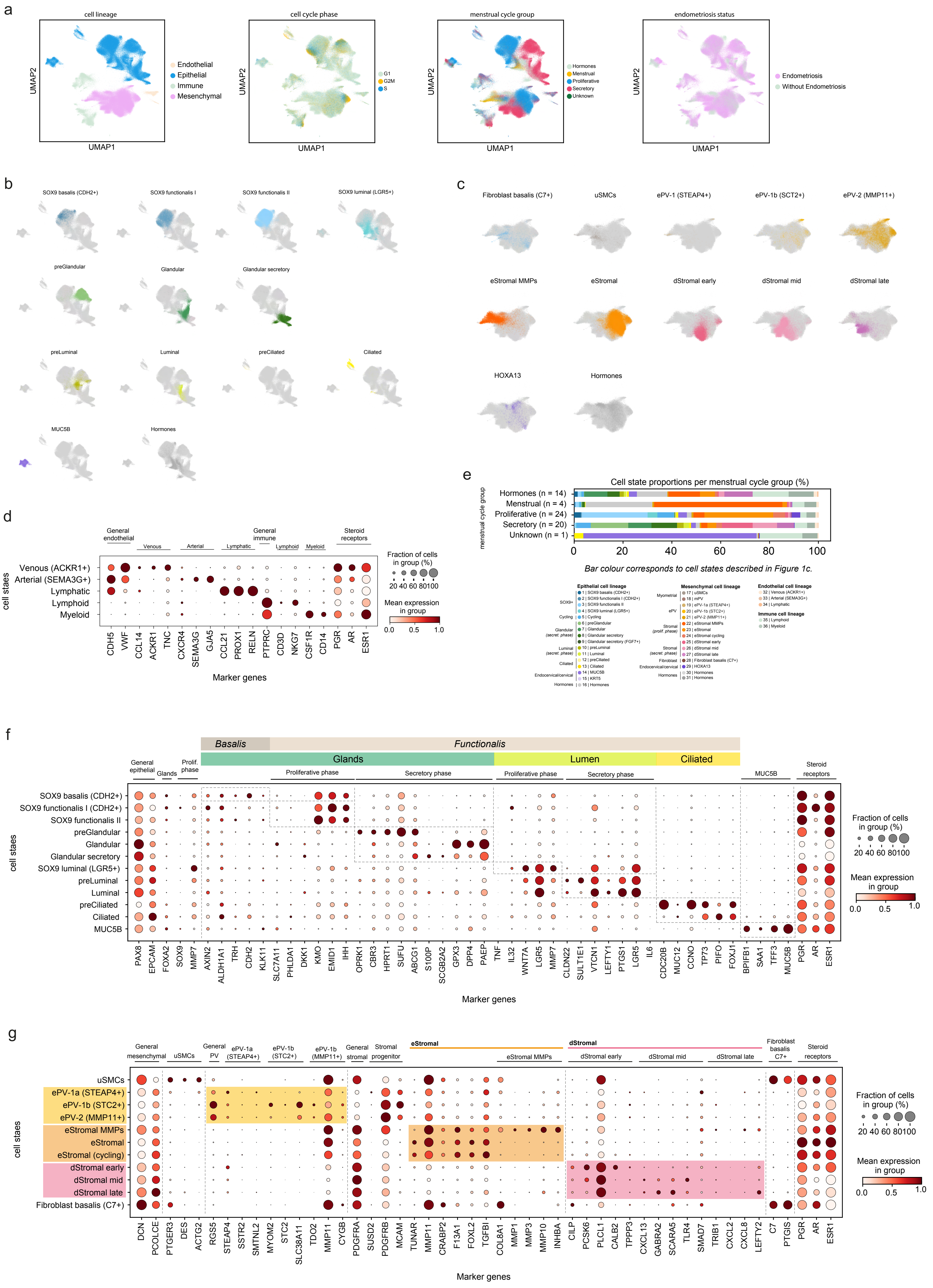

**Supplementary Figure 2. Single-nucleus RNA-sequencing cell state identification and marker gene expression. a,** UMAP projections of the snRNA-seq data coloured by cell lineage, cell cycle phase, menstrual cycle group, and endometriosis status. **b,** UMAP projections of the epithelial cell lineage of the snRNA-seq dataset coloured by the identified epithelial cell states of the HECA as assigned by label transfer. **c,** UMAP projections of the mesenchymal cell lineage of the snRNA-seq dataset coloured by the identified mesenchymal cell states of the HECA as assigned by label transfer. **d,** Dot plot showing normalised, log-transformed and variance-scaled expression of genes (*x*-axis) characteristic of the endothelial and immune nuclei (*y*-axis). **e,** Bar plot showing the cellular composition of endometrial biopsies belonging to the different menstrual cycle groups (*y*-axis). **f,** Dot plot showing normalised, log-transformed and variance-scaled expression of genes (*x*-axis) characteristic of the identified epithelial cell states (*y*-axis) in snRNA-seq data. **g,** Dot plot showing normalised, log-transformed and variance-scaled expression of genes (*x*-axis) characteristic of the identified mesenchymal cell states (*y*-axis) in snRNA-seq data. dStromal, decidualised stromal cells; ePV, endometrial perivascular cells; eStromal, endometrial stromal cells specific to proliferative phase; HECA, human endometrial cell atlas; MMPs, matrix metalloproteinases; mPV, myometrial perivascular cells; Prolif., proliferative; secret., secretory; snRNA-seq, single-nucleus RNA-sequencing; UMAP, uniform manifold approximation and projection; uSMCs, uterine smooth muscle cells.

### Supplementary Figure 3

a

#### Single-cell data

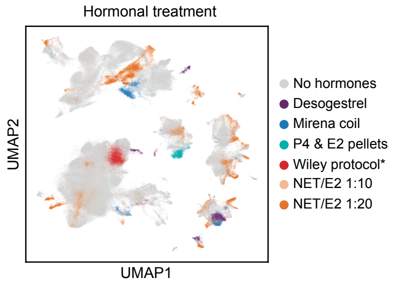


b

| Hormonal treatment | Donor numbers | Cell numbers | Dataset |
|---|---|---|---|
| Desogestrel | 1 | 5,754 | Mareckova |
| Mirena coil | 1 | 6,022 | Mareckova |
| P4 & E2 pellets | 1 | 2,238 | Fonseca |
| Wiley protocol* | 1 | 6,230 | Fonseca |
| NET/E2 1:10 | 2 | 6,331 | Tan |
| NET/E2 1:20 | 8 | 26,376 | Tan |

**E2 and testosterone creams and P4 on days 14-28*
*Abbreviations: E2, oestrogen; NET, norethindrone; P4, progesterone.*

c

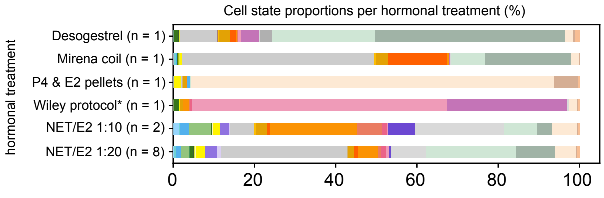


*Bar colour corresponds to cell states below.*

d

#### Single-nuclei data

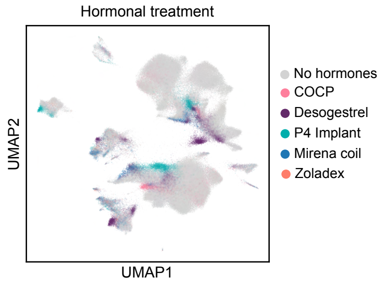


e

| Hormonal treatment | Donor numbers | Nuclei numbers |
|---|---|---|
| COCP | 2 | 8,262 |
| Desogestrel | 4 | 17,035 |
| P4 Implant | 3 | 9,706 |
| Mirena coil | 4 | 6,997 |
| Zoladex | 1 | 2,232 |

*Abbreviations: COCP, combine oral contraceptive pill;*
*P4, progesterone.*

f

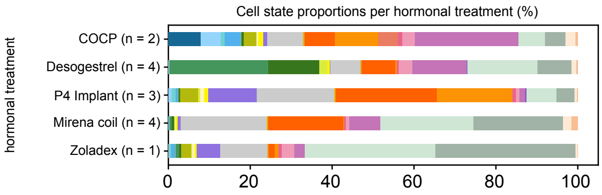


*Bar colour corresponds to cell states below.*

**Epithelial cell lineage**

SOX9+
- 1 | SOX9 basalis (CDH2+)
- 2 | SOX9 functionalis I (CDH2+)
- 3 | SOX9 functionalis II
- 4 | SOX9 luminal (LGR5+)

Cycling
- 5 | Cycling

Glandular *(secret. phase)*
- 6 | preGlandular
- 7 | Glandular
- 8 | Glandular secretory
- 9 | Glandular secretory (FGF7+)

Luminal *(secret. phase)*
- 10 | preLuminal
- 11 | Luminal

Ciliated
- 12 | preCiliated
- 13 | Ciliated

Endocervical/cervical
- 14 | MUC5B
- 15 | KRT5

Hormones
- 16 | Hormones

**Mesenchymal cell lineage**

Myometrial
- 17 | uSMCs
- 18 | mPV

ePV
- 19 | ePV-1a (STEAP4+)
- 20 | ePV-1b (STC2+)
- 21 | ePV-2 (MMP11+)

Stromal *(prolif. phase)*
- 22 | eStromal MMPs
- 23 | eStromal
- 24 | eStromal cycling

Stromal *(secret. phase)*
- 25 | dStromal early
- 26 | dStromal mid
- 27 | dStromal late

Fibroblast
- 28 | Fibroblast basalis (C7+)

Endocervical/cervical
- 29 | HOXA13

Hormones
- 30 | Hormones
- 31 | Hormones

**Endothelial cell lineage**
- 32 | Venous (ACKR1+)
- 33 | Arterial (SEMA3G+)
- 34 | Lymphatic

**Immune cell lineage**
- 35 | Lymphoid
- 36 | Myeloid

**Supplementary Figure 3. Cellular heterogeneity of samples from donors taking exogenous hormones in scRNA-seq and snRNA-seq data.** **a,** UMAP projections of the scRNA-seq data coloured by hormonal treatment taken. **b,** Overview of the number of donors and cells per hormonal treatment taken in each dataset profiled by scRNA-seq. **c,** Bar plot showing the cellular composition of endometrial biopsies from donors taking the different hormonal treatment (*y*-axis) in the scRNA-seq data. **d,** UMAP projections of the snRNA-seq data coloured by hormonal treatment taken. **e,** Overview of the number of donors and cells per hormonal treatment taken profiled by snRNA-seq. **c,** Bar plot showing the cellular composition of endometrial biopsies from donors taking the different hormonal treatment (*y*-axis) in the snRNA-seq dataset. dStromal, decidualised stromal cells; ePV, endometrial perivascular cells; eStromal, endometrial stromal cells specific to proliferative phase; MMPs, matrix metalloproteinases; mPV, myometrial perivascular cells; Prolif., proliferative; scRNA-seq; single-cell RNA-sequencing; secret., secretory; snRNA-seq, single-nucleus RNA-sequencing; UMAP, uniform manifold approximation and projection; uSMCs, uterine smooth muscle cells.

#### Supplementary Figure 4

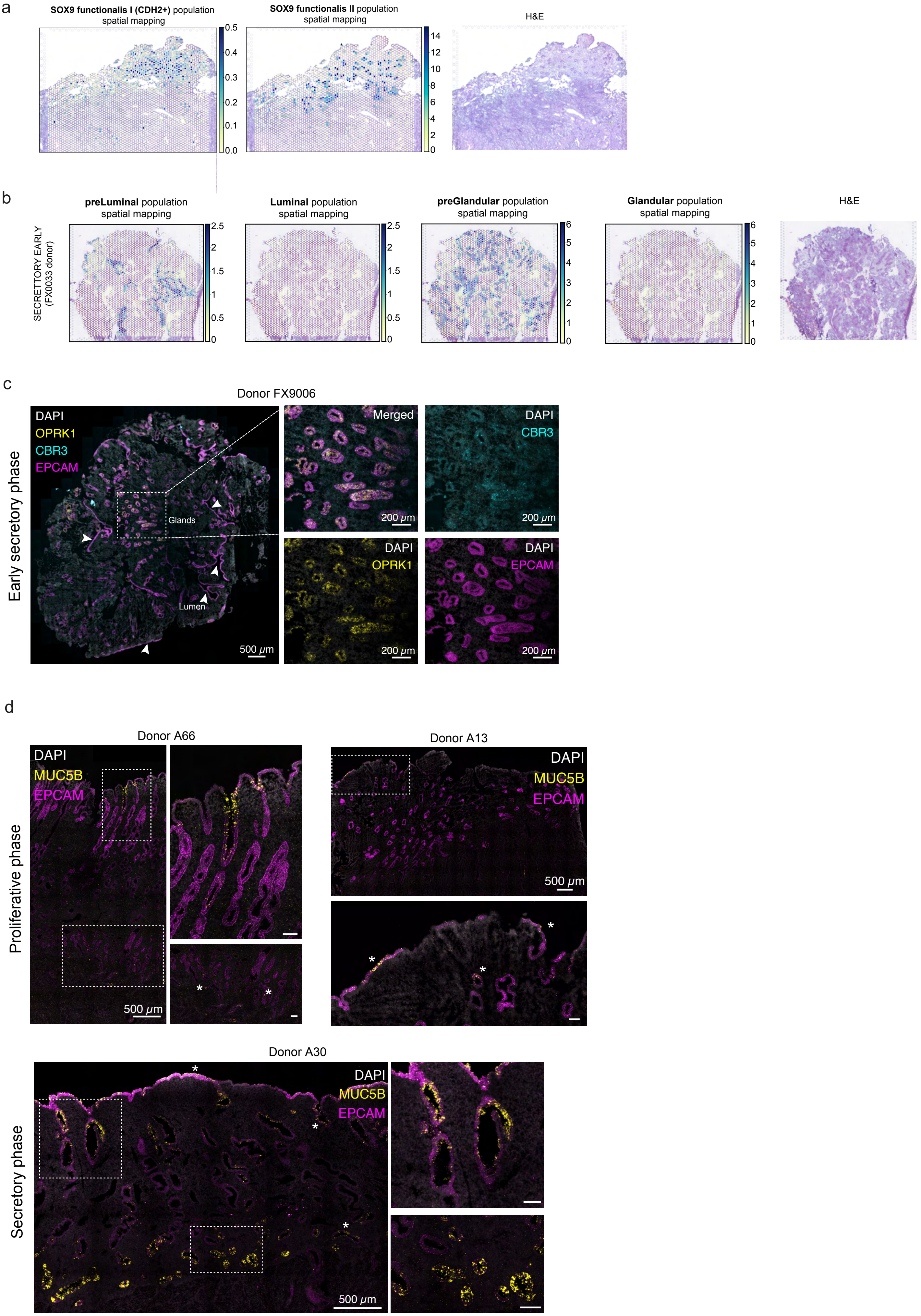

**Supplementary Figure 4. Spatial transcriptomics mapping of epithelial cell populations. a,** Visium spatial transcriptomics data and an H&E image of the same tissue section are shown. Spot colour indicates estimated cell state density for a specific population in each Visium spot, as computed by cell2location. Spatial mapping of the SOX9 functionalis I (CDH2+) and SOX9 functionalis II populations is shown in a section of a whole-uterus biopsy from donor A13. **b,** Visium spatial transcriptomics data and an H&E image of the same tissue section are shown. Spot colour indicates estimated cell state density for a specific population in each Visium spot, as computed by cell2location. Spatial mapping of the preLuminal, Luminal, preGlandular and Glandular populations is visualised in a section of a superficial biopsy from donor FX0033 (early secretory phase). **c,** High-resolution multiplexed smFISH of a section of a superficial biopsy from donor FX9006 (early secretory phase) showing the expression of *DAPI* (white, nuclei), *EPCAM* (magenta, epithelial cells), *CBR3* (cyan, preGlandular cells), and *OPRK1* (yellow, preGlandular cells). The dashed outline indicates the area shown magnified to the right. The magnified image shows the glandular region with high and co-localised *OPRK1* and *CBR3* signal. White arrows indicate luminal regions with low *OPRK1* and *CBR3* signal. **d,** High-resolution multiplexed smFISH of full thickness endometrium sections from the proliferative phase (donors A66 and A13) and secretory phase (donor A30) showing the expression of *DAPI* (white, nuclei), *EPCAM* (magenta, epithelial cells), and *MUC5B* (yellow, epithelial cells). For each panel, the dashed outline indicates the area shown magnified. Asterisks indicate some of the regions where the MUC5B signal was detected and varied across samples. Scale bars are 100 μm, unless differently specified. smFISH, single molecule fluorescence in situ hybridisation.

### Supplementary Figure 5

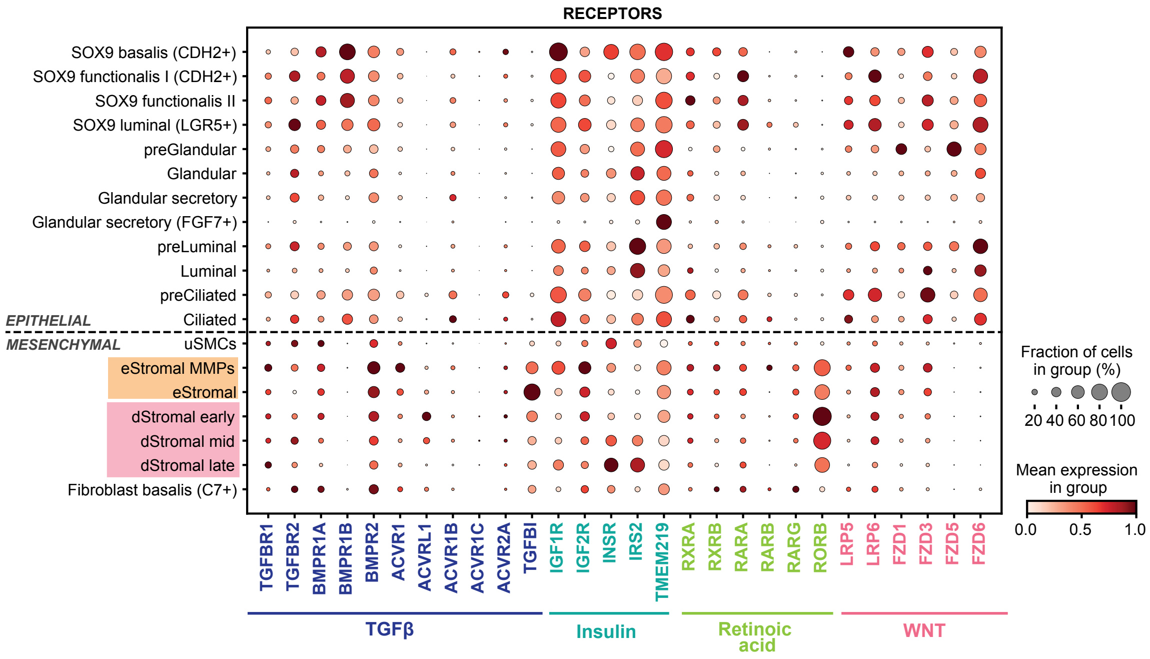

**Supplementary Figure 5. Expression of receptors involved in TGFβ, insulin, retinoic acid and WNT signalling.** Dot plot showing normalised, log-transformed and variance-scaled expression of genes coding for TGFβ, insulin, retinoic acid and WNT signalling receptors (*x*-axis) in the epithelial and mesenchymal cell states identified (*y*-axis) in the scRNA-seq data. eStromal, endometrial stromal cells specific to proliferative phase; dStromal, decidualised stromal cells; MMPs, matrix metalloproteinases; scRNA-seq, single-cell RNA-sequencing; TGFβ, transforming growth factor beta; uSMCs, uterine smooth muscle cells.

#### Supplementary Figure 6

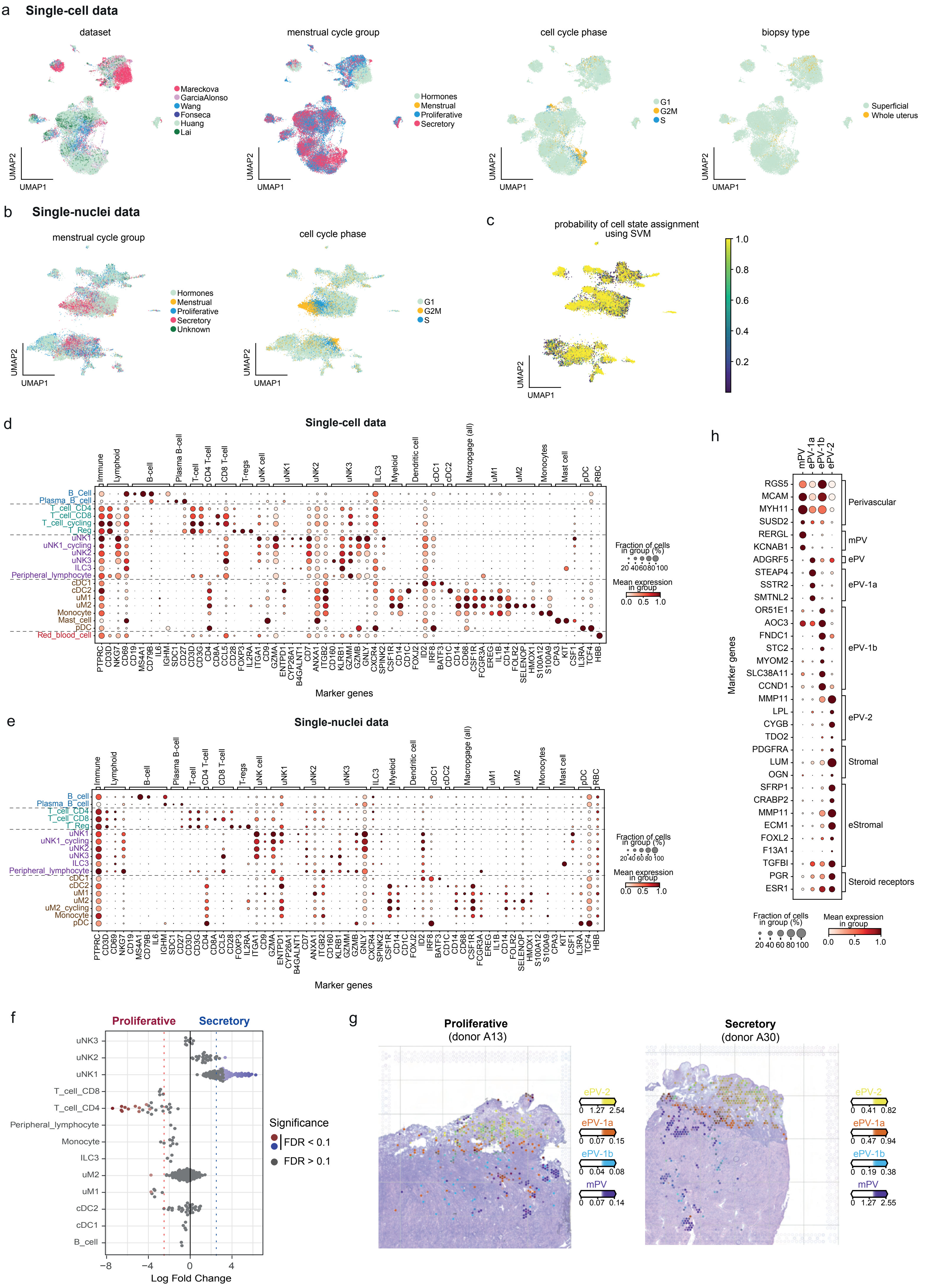

**Supplementary Figure 6. Immune cells in scRNA-seq and snRNA-seq data. a,** UMAP projections of scRNA-seq data for immune cells coloured by dataset, menstrual cycle group, cell cycle phase and biopsy type. **b,** UMAP projections of snRNA-seq data for immune cells coloured by menstrual cycle group and cell cycle phase. **c,** UMAP projection of snRNA-seq data for immune cells coloured by the probability of assigning the immune cell types identified in the scRNA-seq data. Support Vector Machine (SVM) classifier was trained using the immune cell scRNA-seq data and the predicted cell type annotations were then projected onto the snRNA-seq data with the probability shown. **d,** Dot plot showing normalised, log-transformed and variance-scaled expression of genes (*x*-axis) characteristic of the identified immune cell states (*y*-axis) in the scRNA-seq data. **e,** Dot plot showing normalised, log-transformed and variance-scaled expression of genes (*x*-axis) characteristic of the identified immune cell states (*y*-axis) in the snRNA-seq data. **f,** Beeswarm plot of the distribution of log fold change across the menstrual cycle (proliferative and secretory phases) in neighbourhoods containing immune cells from different cell type clusters in snRNA-seq data. Differentially abundant neighbourhoods at log fold change > 2.5 and spatial FDR < 0.1 are coloured. **g,** Visium spatial transcriptomics data for donors A13 (proliferative phase) and A30 (secretory phase) are shown. Spot colour indicates estimated cell state density for a specific population of perivascular cells (mPV, ePV-1a, ePV-1b and ePV-2) in each Visium spot, as computed by cell2location. **h,** Dot plot showing normalised, log-transformed and variance-scaled expression of genes (*x*-axis) characteristic of the identified endothelial, perivascular and stromal cells (*y*-axis) in the scRNA-seq data. cDC, conventional dendritic cells; eStromal, endometrial stromal cells specific to proliferative phase; ePV, endometrial perivascular cells; FDR, false discovery rate; ILC3, innate lymphoid cell type 3; mPV, myometrial perivascular cells; pDC, plasmacytoid dendritic cells; RBC, red blood cells; scRNA-seq, single-cell RNA-sequencing; snRNA-seq, single-nucleus RNA-sequencing; SVM, support vector machine; T Reg, T regulatory cells; uM, uterine macrophages; UMAP, uniform manifold approximation and projection; uNK, uterine natural killer cells.

### Supplementary Figure 7

a

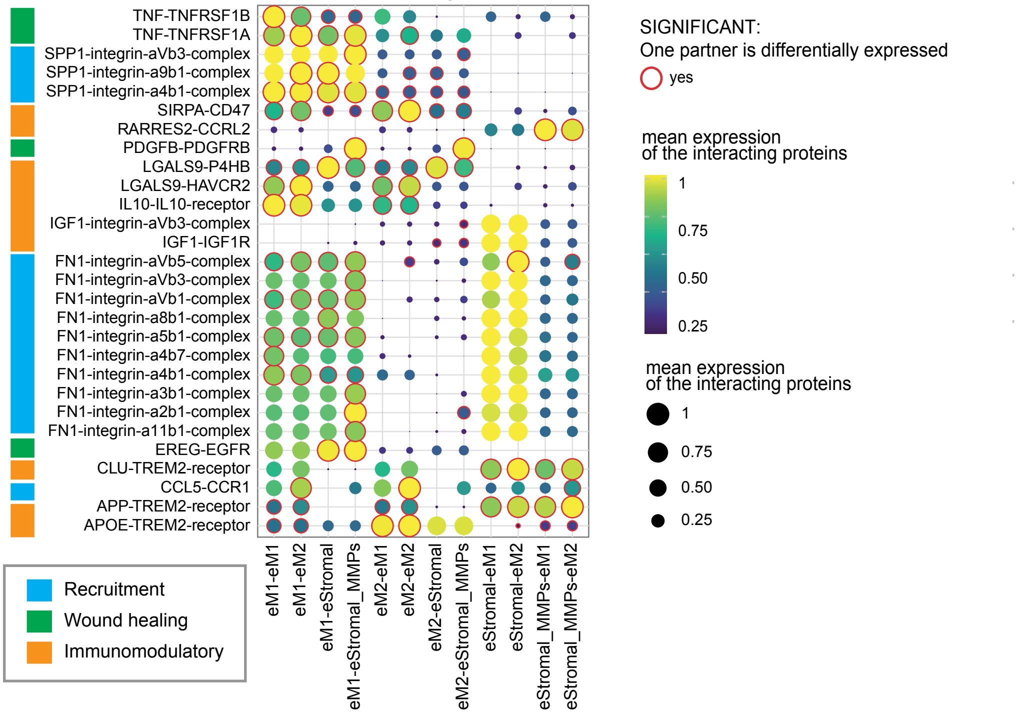


b

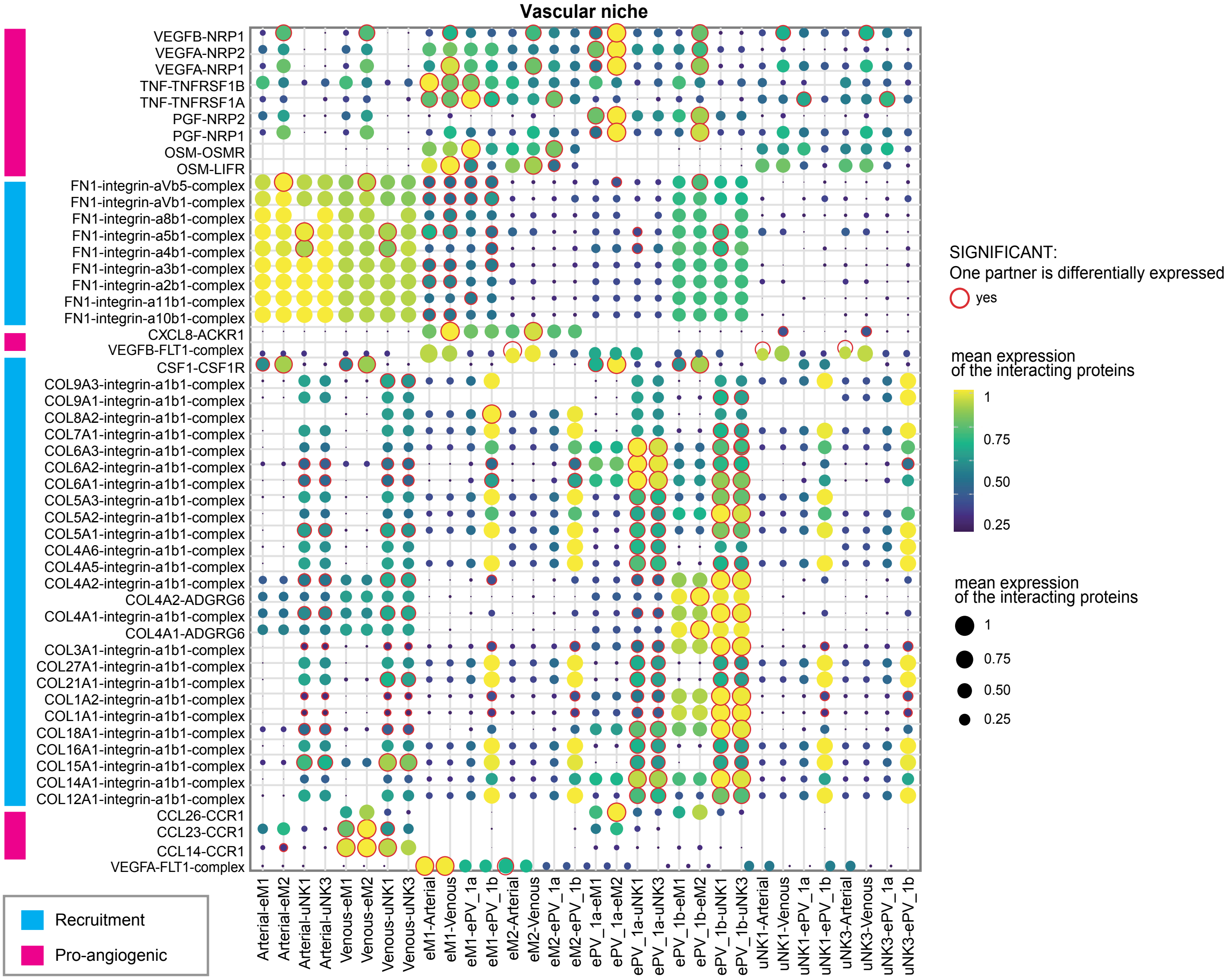

**Supplementary Figure 7. Predicted cell-cell interactions underpinning endometrial regeneration and angiogenesis. a,** Dotplot plot reporting the variance-scaled mean expression of the two or more (if heteromeric complexes) transcripts coding for the interacting proteins in pairs of cell types. Red circles indicate that at least one of the interacting partners is differentially expressed in one of the cell types in the pair. Interactions are classified based on whether they are predicted to play a role in recruitment, wound healing or immunomodulation during endometrial regeneration. **b,** Dotplot plot reporting the variance-scaled mean expression of the two or more (if heteromeric complexes) transcripts coding for the interacting proteins in pairs of cell types. Red circles indicate that at least one of the interacting partners is differentially expressed in one of the cell types in the pair. Interactions are classified based on whether they are predicted to play a role in cell recruitment or pro-angiogenic processes within the vascular niche.
